# Dysregulation of xenobiotic metabolism and mitochondrial dysfunction exacerbate acetaminophen-induced hepatotoxicity in human antigen R-deficient male mice

**DOI:** 10.64898/2026.01.28.702297

**Authors:** Natalie Eppler, Elizabeth Jones, Forkan Ahamed, Naren Raja, Jephte Y. Akakpo, Margitta Lebofsky, Lily He, Ira Vats, Priyanka Ghosh, Yifan Yu, Kaitlyn Thomas, Colin McCoin, John Thyfault, Xiaoqing Wu, Liang Xu, Wei Cui, Rong Wang, Hartmut Jaeschke, Yuxia Zhang

**Affiliations:** Department of Pharmacology, Toxicology and Therapeutics, University of Kansas Medical Center, Kansas 66160, USA; Department of Oral and Craniofacial Sciences, School of Dentistry, University of Missouri-Kansas City, Missouri 64108, USA; Departments of Cell Biology and Physiology, KU Diabetes Institute, and Kansas Center for Metabolism and Obesity Research, University of Kansas Medical Center, Kansas 66160, USA; Department of Molecular Biosciences, University of Kansas, Lawrence, Kansas 66045, USA; Department of Pathology, University of Kansas Medical Center, Kansas 66160, USA

**Keywords:** Acetaminophen hepatotoxicity, Human antigen R, gene knockout, male mouse, xenobiotic metabolism, mitochondrial function

## Abstract

Acetaminophen (APAP) overdose is a leading cause of acute liver failure worldwide. The RNA-binding protein Human antigen R (HuR) is a multifunctional post-transcriptional regulator that plays a pivotal role in cellular stress responses, including those triggered by APAP toxicity. This study investigated the mechanisms by which HuR protects against APAP-induced hepatotoxicity in male mice. Hepatocyte-specific *HuR*-deficient (*HuR*^Hep-/-^) male mice on a C57BL/6N background and wild-type (WT) littermates were treated with 200 mg/kg APAP, and liver tissues were collected at 2, 6, and 24 hours post-treatment. APAP administration increased hepatic *HuR* mRNA expression and induced HuR cleavage and the formation of a higher-molecular weight HuR-immunoreactive band, with the latter two correlating with injury severity. Compared with WT controls, *HuR*^Hep-/-^ mice exhibited markedly increased susceptibility to hepatotoxicity at both 2 and 6 hours. Metabolite profiling revealed altered APAP metabolism and reduced glutathione S-transferase (Gst) expression in *HuR*^Hep-/-^ livers, consistent with impaired APAP detoxification and increased APAP-protein adduct formation. Fourier-transform infrared (FTIR) spectroscopy further identified early biochemical differences between WT and *HuR*^Hep-/-^ livers as early as 2 hours after APAP exposure. Additionally, *HuR* deficiency resulted in pronounced mitochondrial structural abnormalities and dysfunction at 2 and 6 hours, accompanied by reduced expression of the mitochondrial fission and fusion proteins Drp1 and Mfn2, increased mitochondrial protein release, and enhanced hepatocyte death. Although pro-inflammatory cytokine levels were elevated in *HuR*^Hep-/-^ mice relative to WT controls at 24 hours, hepatocyte proliferation was similarly blunted in both genotypes, consistent with severe liver injury and delayed recovery. Collectively, these findings identify hepatocyte HuR as a critical regulator of xenobiotic metabolism and mitochondrial integrity and establish its essential role in early protection against APAP-induced hepatotoxicity in male mice.

## INTRODUCTION

Globally, acetaminophen (APAP), also known as paracetamol, is among the most widely used analgesic and antipyretic medications (Chidiac et al. 2023). Due to its narrow therapeutic index and widespread presence in over-the-counter medications, APAP is the leading cause of acute liver failure in the United Kingdom and the United States (Bernal and Wendon 2013). Early administration of the sulfhydryl donor, N-acetylcysteine (NAC), is key to detoxification and patient survival following an APAP overdose (Green et al. 2013). However, APAP overdose patients often present late to the clinic, limiting the efficacy of current intervention strategies (Chidiac et al. 2023). To improve current intervention strategies, a greater mechanistic understanding of APAP-induced hepatotoxicity is warranted.

The liver plays a central role in the metabolism and elimination of APAP. At therapeutic doses, APAP is primarily detoxified through glucuronidation and sulfation to form non-reactive metabolites (McGill and Jaeschke 2013). However, at supratherapeutic doses, the sulfation pathway becomes saturated, leading to increased formation of the hepatotoxic metabolite N-acetyl-p-benzoquinone imine (NAPQI) (Ramachandran and Jaeschke 2019a). Rapid detoxification of NAPQI through conjugation with glutathione (GSH) is essential to prevent hepatic injury (Mitchell et al. 1973). The resulting APAP conjugates are subsequently eliminated from hepatocytes via apical and basolateral membrane transport proteins (McGill and Jaeschke 2013). Experimental studies targeting various drug-metabolizing enzymes (DMEs) and membrane transporters have revealed complex and overlapping mechanisms governing APAP detoxification and disposition (Henderson et al. 2000; Zamek-Gliszczynski et al. 2005; Zamek-Gliszczynski et al. 2006). Although the number of DMEs is relatively limited compared with the vast array of xenobiotics they process, modulation of their expression through induction or repression has profound effects on xenobiotic half-life, activation, and tissue distribution (Parkinson et al., 2018). Nuclear receptors such as pregnane X receptor (PXR) and constitutive androstane receptor (CAR) are well established transcriptional regulators of DMEs in this context (Willson and Kliewer 2002). In contrast, the post-transcriptional regulation of DME expression remains poorly understood. Tight control of mRNA stability is a fundamental mechanism that enables cells to rapidly adapt to changing environmental conditions. Accordingly, a major objective of this study was to elucidate the role of the RNA-binding protein Human antigen R (HuR) in the post-transcriptional regulation of hepatic APAP metabolism.

Mitochondria are particularly susceptible to NAPQI-mediated toxicity, and interventions that mitigate mitochondrial injury provide significant protection against APAP-induced hepatotoxicity (Ramachandran et al. 2018). Within mitochondria, covalent binding of NAPQI to nucleophilic proteins disrupts the respiratory chain, enhances reactive oxygen species generation, dissipates mitochondrial membrane potential, and ultimately triggers opening of the mitochondrial permeability transition (MPT) pore, leading to necrotic cell death (Ramachandran and Jaeschke 2019b). Moreover, disturbances in mitochondrial morphology and dynamics impair respiratory efficiency (Donnelly et al. 1994) and increase mitochondrial susceptibility to MPT pore opening (Picard et al. 2013). In the liver, HuR is activated in response to oxidative stress and confers cyto-protection in murine hepatocytes during ischemia reperfusion injury, in part through induction of the antioxidant enzyme heme oxygenase-1 (HO-1) (Dery et al. 2020). Similarly, HuR protects against diet-induced mitochondrial dysfunction by regulating expression of key electron transport chain components, including cytochrome c (Cytc), NADH:ubiquinone oxidoreductase subunit B6 (Ndufb6), and ubiquinol–cytochrome c reductase binding protein (Uqcrb) (Zhang et al. 2020). Recent evidence also indicates that hepatocyte HuR protects against APAP-induced liver injury by promoting hepatocyte proliferation, autophagy, and antioxidant defense following overdose (Lu et al. 2024). However, whether the exacerbated liver injury observed in hepatocyte-specific *HuR*-deficient (*HuR*^Hep-/-^) mice results, at least in part, from mitochondrial dysfunction and increased oxidative stress remains unclear. Therefore, the second aim of this study was to evaluate mitochondrial structural and functional integrity in *HuR*^Hep-/-^ livers under normal condition and over the time course following APAP overdose.

Fourier transform infrared (FTIR) spectroscopy is a label-free and highly sensitive technique for detecting subtle biomolecular alterations in biological samples (Baker et al. 2014). By measuring infrared light absorption, FTIR generates unique molecular fingerprints, particularly within the fingerprint region (1800–900 cm⁻¹), which corresponds to vibrational transitions of proteins, lipids, carbohydrates, and nucleic acids (Wang and Wang 2021). Our previous studies demonstrated that FTIR-based biomolecular profiling can effectively differentiate oral squamous cell carcinoma from benign tissues (Wang et al. 2021), predict malignant transformation risk in precancerous oral lesions (Wang et al. 2025), and identify biochemical signatures associated with tumor treatment responses (Ly et al. 2025). Despite its proven utility in oncology, the application of FTIR spectroscopy in detecting early biochemical alterations in liver tissues following drug-induced injury remains unexplored. Therefore, the third aim of this study was to employ FTIR spectroscopy to characterize subtle molecular differences between wild-type (WT) and *HuR*^Hep-/-^livers following APAP overdose, thereby assessing its potential as a sensitive, non-destructive analytical tool for early detection of molecular perturbations in liver toxicology research.

In the present study, we demonstrate that the disposition of GSH-conjugates of APAP metabolites is markedly altered in *HuR*^Hep-/-^ mice, a change that is associated with dysregulation of glutathione S-transferase (Gst), the phase II DMEs. Consistent with these findings, FTIR analysis revealed altered hepatic composition in *HuR*-deficient mouse livers following APAP overdose. In addition, *HuR*^Hep-/-^ livers consistently exhibited reduced mitochondrial abundance and disrupted mitochondrial morphology, accompanied by reduced expression of the mitochondrial fusion and fission proteins Mfn2 and Drp1, impaired respiratory capacity, and increased ROS production. Together, these data support a protective role for HuR in maintaining liver function during acute APAP-induced injury and reveal multiple regulatory mechanisms through which HuR exerts these effects. Collectively, our findings provide new insight into the molecular networks governed by hepatocyte HuR, highlighting its essential role in xenobiotic metabolism, mitochondrial integrity, and protection against APAP-induced hepatotoxicity.

## MATERIALS AND METHODS

### Animal Studies

*HuR*^flox/flox^ mice (JAX stock #: 021431) and Alb-Cre mice (JAX stock #: 003574) were obtained from Jackson Laboratory (Bar Harbor, ME, USA). Hepatocyte *HuR* knockout (*HuR*^Hep-/-^represents HuR^flox/flox^; Alb-Cre positive) and their littermate wild-type controls (WT represents HuR^flox/flox^; Alb-Cre negative) were generated by crossbreeding *HuR*^flox/flox^ mice with Alb-Cre mice and subsequently backcrossing into the C57BL/6N genetic background for 10 generations. Mice were housed in a virus-free facility with a 12 h light/dark cycle (lights on from 6 a.m. to 6 p.m.) and maintained at a temperature of 25°C, with ad libitum access to food and water. Male mice aged 8-10 weeks were used for the experiments (n=5-6/group), unless otherwise stated. Prior to APAP administration, mice were fasted overnight. APAP from Sigma-Aldrich (St. Louis, MO, USA) was dissolved in warm saline and administered intraperitoneally at a dose of 200 mg/kg. All experiments were conducted in compliance with relevant guidelines and regulations approved by the Institutional Animal Care and Use Committee (ICAUC) at the University of Kansas Medical Center.

### Histopathology and Immunohistochemistry

Liver tissues were fixed in 10% formalin in PBS (pH 7.4) for 24 hours, then processed, embedded in paraffin, sectioned into 5 μm slices, and stained with hematoxylin and eosin (H&E) to assess histopathology. Terminal deoxynucleotidyl transferase dUTP nick end labeling (TUNEL) staining was performed on rehydrated liver sections using the In Situ Cell Death Detection Kit (Roche, 11684809910) to assess cell death. For immunohistochemistry (IHC), antigen retrieval was performed on rehydrated sections and sodium citrate buffer (10 mM, pH 6.0) in a pressure cooker at high pressure for 5 minutes was used for F4/80, while Tris-EDTA buffer (10 mM, pH 9.0) at high pressure for 10 minutes was used for PCNA and Ly6g. Endogenous peroxidase activity was blocked with 0.3% H₂O₂, and non-specific binding was prevented with 10% normal horse serum. Sections were incubated overnight at 4°C with primary antibodies: F4/80 (Cell Signaling Technology, 70076), PCNA (Cell Signaling Technology, 13110S), and Ly6g (Cell Signaling Technology, 87048). Antibody binding was detected using ImmPRESS peroxidase polymer detection kits (Vector Laboratories, MP-7402) and ImmPACT DAB substrate (Vector Laboratories, SK-4105). Sections were counterstained with hematoxylin. An anti-3-Nitrotyrosine antibody (Abcam, ab61392) was used for immunofluorescence staining. Images were then acquired with an Olympus IX73 microscope and Fiji (ImageJ) was used for quantification.

### FTIR imaging and data analysis

Paraffin-embedded liver tissues sectioned at 5 µm thickness were mounted on barium fluoride (BaF₂) discs (25 × 2 mm; REFLEX Analytical Corp., Ridgewood, NJ, USA) for FTIR imaging. Sections were deparaffinized in histological-grade xylene (CAS 1330-20-7; Sigma-Aldrich, St. Louis, MO, USA) for 5 min × 3 at room temperature, air-dried, and stored overnight in a vacuum desiccator. FTIR images were acquired in transmission mode using a PerkinElmer Spectrum Spotlight imaging system (Spectrum One, Spotlight 300; PerkinElmer, Waltham, MA, USA) with Spectrum IMAGE software. Imaging parameters were as follows: spectral resolution, 4 cm⁻¹; spectral range, 4000–950 cm⁻¹; pixel size, 6.25 µm; and 16 co-added scans per pixel. Three regions of interest (ROIs), each measuring 400 × 400 µm and located near central vein regions, were selected per tissue section. Spectral preprocessing and analysis were conducted using PLS_Toolbox (Eigenvector Research Inc., Manson, WA, USA) in MATLAB R2020b (MathWorks, Natick, MA, USA). The workflow included: (1) transmission-to-absorbance conversion (A = log(1/T)); (2) selection of the fingerprint region (1800–950 cm⁻¹); (3) Savitzky–Golay smoothing; (4) extended multiplicative signal correction; (5) automated weighted least-squares baseline correction; and (6) vector normalization. Principal component analysis (PCA) and Hotelling’s T² versus Q-residuals plots were then applied for exploratory analysis and outlier removal (Wang et al. 2021). A representative FTIR spectrum for each ROI was obtained by averaging pixel-level spectra and used for subsequent analysis.

### Determination of APAP metabolites in mouse plasma by LC-MS/MS

APAP metabolites in mouse plasma were detected as described previously (Akakpo et al. 2018; Ghosh et al. 2023). Standards, including 4-acetamidophenyl β-D-glucuronide (APAP-Gluc), 4-acetaminophen sulfate (APAP-Sulf), 3-(N-acetylcysteine) acetaminophen (APAP-NAC), 3-cysteinylacetaminophen trifluoroacetic acid (APAP-Cys), acetaminophen glutathione (APAP-GSH), along with internal standards acetaminophen-d4 and acetaminophen-sulfate-d3, were sourced from Toronto Research Chemicals (Toronto, Canada). Stock solutions (1 mM) and a mixed standard solution (75 μM) were prepared in 50:50 methanol: water, with working standards (0.25–25 μM) diluted from the stock. Samples and standards were prepared on ice. Plasma from untreated mice was pooled as the blank matrix. To precipitate proteins, 20 μl of plasma or 50 μl of working standard was mixed with 90 μl of methanol containing internal standards (200 ng/ml acetaminophen-d4 and 1000 ng/ml acetaminophen-sulfate-d3). Additional water and methanol were added, followed by vortexing for 10 seconds and centrifugation at 13,400 × g for 10 minutes at 4°C. Analysis was performed using LC-MS/MS with a Waters Acquity UPLC® system and Quattro Premier XE mass spectrometer in positive mode with multiple reaction monitoring (MRM). Separation was achieved on a Waters UPLC® HSS T3 column (1.8 μm, 2.1 × 150 mm) at 50°C using a gradient flow of mobile phase A (6 mM ammonium acetate in water with 0.01% formic acid) and mobile phase B (methanol). The gradient transitioned from 2% to 75% B over 3.5 minutes, then to 98% B over 0.5 minutes, and held for 2 minutes at 0.4 ml/min. Quantification used a weighted (1/x) linear regression of analyte/internal standard peak area ratios. Internal standard acetaminophen sulfate-d3 and QuanLynx software (version 4.1) were used for MS peak integration. Detection limits were 0.250 μM for APAP-GSH, 0.125 μM for APAP-Sulf and APAP-Gluc, 0.063 μM for APAP-Cys, and 0.025 μM for APAP-NAC.

### Analysis of APAP protein adducts by high-pressure liquid chromatography (HPLC)

Liver APAP protein adducts were quantified as described previously (Akakpo et al. 2018; Ghosh et al. 2023). Briefly, Liver homogenates were filtered through a Bio-Spin 6 column to remove lower molecular weight metabolites. To release APAP-Cys, filtrates were then digested overnight with 8 U/ml Streptomyces griseus protease. Following digestion, lysates were precipitated using 40% trichloroacetic acid for 10 minutes on ice. The residue of APAP-Cys adducts was analyzed using HPLC with an electrochemical detector (HPLC-ECD).

### Hepatic mitochondrial isolation and measurement of respiration and H_2_O_2_ emission

Hepatic mitochondria were isolated as described previously (Kumari et al. 2024; McCoin et al. 2019). Briefly, liver tissue was excised, submerged in 8 mL cold isolation buffer (220 mM mannitol, 70 mM sucrose, 10 mM Tris, 1 mM EDTA, pH 7.4), and homogenized on ice. Homogenates were centrifuged at 4°C for 10 minutes at 1500 × g. The supernatant was strained and centrifuged again at 4°C for 10 minutes at 8000 × g. The mitochondrial pellet was resuspended in 6 mL isolation buffer and centrifuged at 4°C for 10 minutes at 6000 × g. This step was repeated using 4 mL isolation buffer containing 0.1% fatty acid-free BSA. The final pellet was resuspended in MiR05 mitochondrial respiration buffer (pH 7.1) and used for respiration studies. Mitochondrial respiration and H_2_O_2_ emission were measured simultaneously using an Oroboros O2K fluorometer (Oroboros Instruments, Austria) following established protocols (Kumari et al. 2024; McCoin et al. 2019). Experiments started with 2 mM malate, 10 mM coenzyme A, and 2.5 mM L-carnitine. Potassium pyruvate (5 mM) was added to measure basal respiration (Leak). Maximal coupled respiration (State 3) was determined with 2.5 mM adenosine diphosphate (ADP) and further complex I-driven rates were measured with 2 mM glutamate. Maximal respiratory rates (State 3S) were assessed by adding 10 mM succinate, and uncoupled respiration was measured with 0.1 mM carbonyl cyanide-p-(trifluoromethoxy) phenylhydrazone (FCCP) titrations. All data were normalized to mitochondrial protein content determined by a DC protein assay (BIO-RAD, Hercules, CA). OXPHOS capacity was calculated as State 3 respiration minus Leak, and mitochondrial redox was determined as basal H_2_O_2_ production divided by basal respiration. Data were analyzed using DatLab 7 Software.

### Electron Microscopy

Liver pieces (1 x 5 mm) were fixed with 2% glutaraldehyde in 0.1 M cacodylate buffer for 2 hours at room temperature with shaking, followed by overnight fixation at 4 °C. Tissues were then washed in 1% OsO_4_ and embedded in Embed 812 resin. Ultrathin sections were positioned within grids and stained with uranyl acetate and lead citrate. Sections were viewed on a JEOL JEM-1400 transmission electron microscope equipped with a Lab6 gun (JEOL, Tokyo, Japan). Images were acquired digitally, and mitochondrial features were assessed using Fiji (ImageJ) software.

### Biochemical measurements of GSH/GSSG and alanine aminotransferase (ALT)

Total liver GSH levels were determined using a modified Tietze assay (Jaeschke and Mitchell 1990). In brief, snap-frozen livers were homogenized on ice in 3% sulfosalicylic acid containing 0.1 mM EDTA. To measure liver GSH, one aliquot of liver homogenate was added to 0.01 N HCl, centrifuged and the supernatant was further diluted with 100mM potassium phosphate buffer (KPP). To measure GSSG, another aliquot was added to 10mM N-ethylmaleimide (NEM) in KPP to trap GSH. The residual NEM was removed with a C18 SepPack column and GSSG was determined by the Tietze assay using dithionitrobenzoic acid. Plasma ALT was measured spectrophotometrically with an ALT reagent kit according to the manufacturer’s instructions (Pointe Scientific, Canton, MI).

### RNA isolation and real-time PCR

Total RNA was isolated using TRIzol reagent (Invitrogen), as described previously (Magee et al. 2020; Zou et al. 2018). cDNA was synthesized from Isolated RNA using the Applied Biosystems High-Capacity cDNA Reverse Transcription Kit (Fisher Scientific, Hampton, NH, USA). Real-time PCR was performed using Applied Biosystems SYBR Green PCR Master Mix (Fisher Scientific, Hampton, NH, USA). The primer sequences used for PCR amplification can be found in Supplementary Table 1. The cDNAs from 3 to 5 animals within each group were pooled and subjected to triplicate qPCR analyses. The amplification results were quantified by measuring the threshold cycle (Ct) value and normalized to the expression of the housekeeping genes glyceraldehyde-3-phosphate dehydrogenase (*Gapdh*) or hypoxanthine guanine phosphoribosyl transferase 1 (*Hprt1*). The data were presented as the fold change in the tested group compared with the control group.

### RNA sequencing data analysis

Liver transcriptome profiles from *HuR*^Hep-/-^ and WT fed a normal chow diet were analyzed using the RNA sequencing (RNA-seq) data set GSE287727, recently deposited to GEO by our group. Analysis was conducted with CLC Genomics Workbench software (Qiagen, Redwood City, CA, USA). Illumina reads were aligned to the *Mus musculus* C57BL/6 genome refseq_GRCm38.p6 using local read and reverse strand alignment. Reads were visualized, and differentially expressed genes (DEGs) were identified based on an adjusted p-value of ≤0.05 and a fold change ≥ 1.5 relative to controls. DEGs with an RPKM value < 0.01 (∼10 total counts) in either of the *HuR*^Hep-/-^ and WT groups were excluded from further analysis. Canonical pathway enrichment was determined using Ingenuity Pathway Analysis (IPA) software (Qiagen, Redwood City, CA, USA), with pathways considered significant if they had a z-score ≥ 2.

### Protein isolation and immunoblotting

Mouse liver protein lysates were prepared by homogenizing tissues in RIPA buffer (20 mM Tris-HCl (pH 7.5), 150 mM NaCl, 1 mM Na2EDTA, 1 mM EGTA, 1% NP-40, 1% sodium deoxycholate, 2.5 mM sodium pyrophosphate, 1 mM beta-glycerophosphate, 1 mM Na_3_VO_4_, 1 µg/ml leupeptin) with protease inhibitors (Fisher Scientific PI78410, Hampton, NH, USA). After centrifugation the protein-containing supernatant was collected for whole protein analysis. Cytoplasmic and mitochondrial protein fractions were isolated as described (Adelusi et al. 2022). Liver tissue was homogenized in ice-cold mitochondrial isolation buffer (220 mM mannitol, 70 mM sucrose, 2.5 mM HEPES, 10 mM EDTA, 1 mM EGTA, 0.1% BSA, pH 7.4) using a Teflon pestle. Homogenates were centrifuged at 2,500 g for 10 minutes at 4°C to remove nuclei and debris. The supernatant was centrifuged again at 20,000 g for 10 minutes at 4°C. The cytoplasmic protein was collected from the supernatant, and the mitochondrial pellet was washed and centrifuged again. Protein concentrations were measured using the DC protein assay (BIO-RAD, Hercules, CA, USA). Protein samples were mixed with loading buffer and resolved on polyacrylamide gels by electrophoresis and transferred to nitrocellulose membranes. Membranes were blocked with 5% skim milk for 1 hour at room temperature, then incubated with primary antibodies (1:1000 dilution) either overnight at 4°C or for 2 hours at room temperature. Primary antibodies included rabbit anti-AIF (CST, 4642S), rabbit anti-Bax (CST, 2772S), rabbit anti-Smac (CST, D5S3R), mouse anti-HuR (Santa Cruz, sc-5261), rabbit anti-Cyp2e1 (Abcam, Ab28146), rabbit anti-pJNK (CST, 4668P), rabbit anti-JNK (CST, 9252T), rabbit anti-Caspase-3 (CST, 9665), rabbit anti-cyclin D1 (CST, 2978S), mouse anti-GAPDH (Sigma, G8795), mouse anti-α-tubulin (Sigma, T6199), mouse anti-β-actin (Sigma, A2228), and rabbit anti-VDAC (CST, D73D12). Following primary antibody incubation, membranes were washed and incubated with the corresponding horseradish peroxidase-conjugated secondary antibody at room temperature for 1 h. Antibody binding was visualized using either Super Signal West Pico Plus Chemiluminescent Substrate (Fisher, PI34580) or Super Signal West Femto Chemiluminescent Substrate (Fisher, PI34094). Images were captured using an Odyssey XF imaging system (LI-COR, Lincoln, NE). β-Actin, GAPDH, and VDAC were used as loading controls, and protein band densitometry was performed using Image Studio Lite software. Expression levels were normalized to the loading controls.

### Statistical Analysis

GraphPad Prism 9.0 (GraphPad Software, La Jolla, CA, USA) was used for data analysis. Data are expressed as mean ± SEM (standard error of the mean). Statistical differences between 2 groups were analyzed using an unpaired 2-tailed Student’s t test. A p value of less than 0.05 was considered statistically significant.

## RESULTS

### APAP overdose increases hepatic *HuR* mRNA expression and induces HuR cleavage and formation of a higher-molecular weight HuR-immunoreactive product

Cellular stress is known to alter HuR expression, subcellular localization, post-translational protein modification, and protein proteolytic cleavage (Mazroui et al. 2008; Talwar et al. 2011). In this study, hepatic HuR expression was assessed over the time course of APAP-induced hepatotoxicity in C57BL/6N male mice treated with 200 mg/kg APAP. Increases in plasma ALT measured at 2, 6, and 24 hours post-APAP overdose were consistent with previous reports in C57BL/6J mice (Bhushan et al. 2014; Duan et al. 2016) (Figure 1A). Rising plasma ALT levels also coincided with increased liver *HuR* mRNA expression at 2 and 6 hours following APAP treatment, whereas *HuR* mRNA levels returned to baseline by 24 hours (Figure 1B).

**Figure 1.**
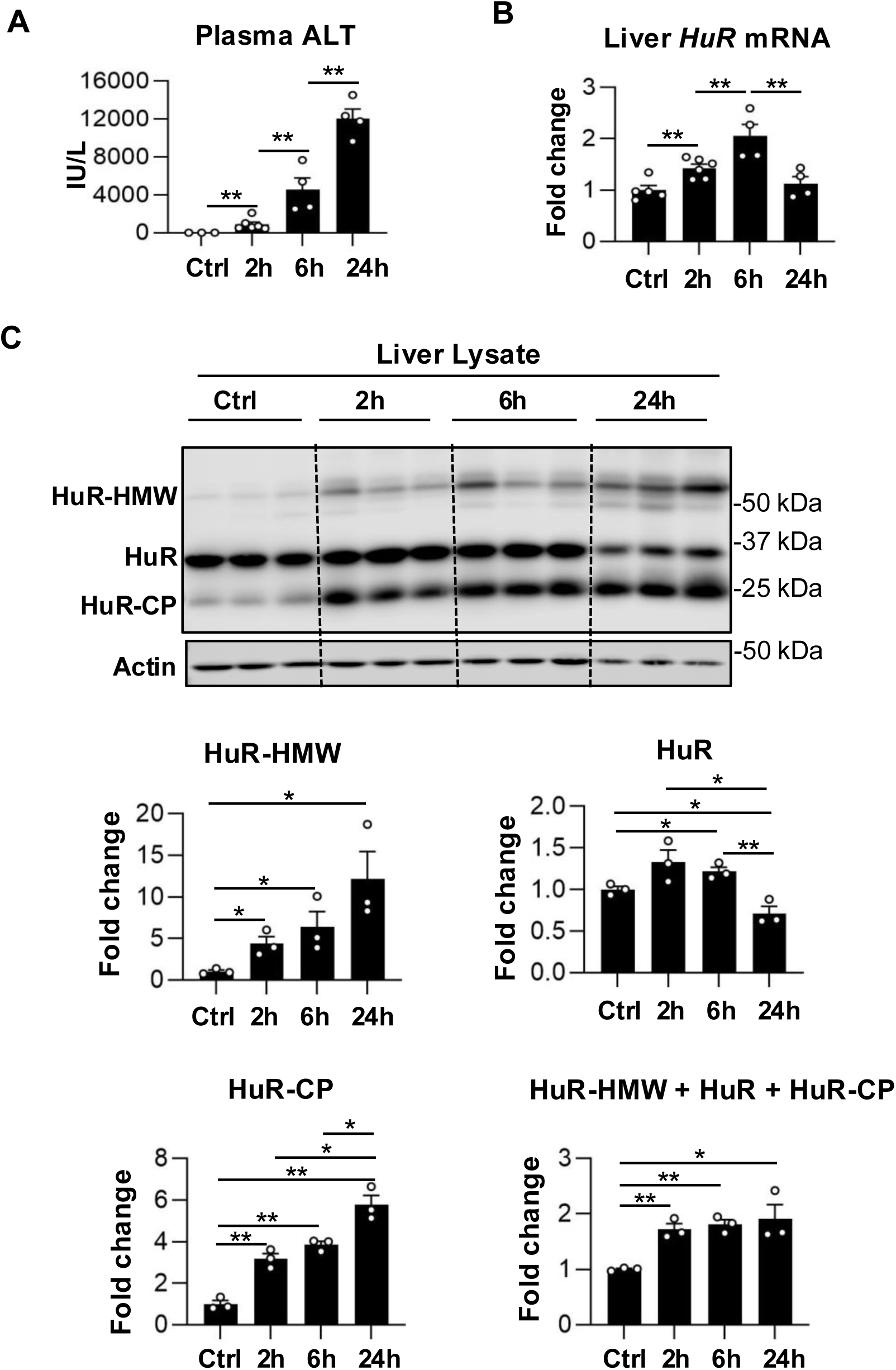
Acetaminophen overdose increases hepatic HuR mRNA expression and induces HuR cleavage and formation of a higher-molecular weight HuR-immunoreactive band in male mice. Measurements were obtained from liver and plasma samples collected at 2, 6, and 24 hours following a 200 mg/kg APAP overdose. (A) Plasma alanine aminotransferase (ALT) levels following APAP overdose. (B) qPCR analysis of hepatic HuR mRNA expression. (C) Western blot analysis of HuR in whole liver lysates revealed three HuR-immunoreactive bands: a higher–molecular weight species (HuR-HMW), full-length HuR (32 kDa), and a HuR cleavage product (HuR-CP, 24 kDa). Band intensities were quantified and normalized to the loading control β-actin. Data are presented as mean ± SEM. n=3-5 mice per group. *p < 0.05, **p < 0.01. Abbreviations: APAP, acetaminophen; WT, wild type; ALT, aminotransferase.

Western blot analysis of liver samples following APAP exposure revealed the appearance of two additional HuR-immunoreactive bands beyond the full-length protein: a higher-molecular weight band (HuR-HMW) and a lower-molecular weight cleavage product (HuR-CP) (Figure 1C). Both HuR-HMW and HuR-CP were minimally detectable in untreated control livers but increased markedly after APAP exposure, with their abundance closely paralleling serum ALT levels. Lethal cellular stress has been known to trigger caspase-mediated cleavage of HuR at aspartate 226 within its nucleocytoplasmic shuttling sequence, generating the 24-kDa HuR cleavage product (HuR-CP) that contributes to pp32/PHAP-I-mediated regulation of apoptosis (Mazroui et al. 2008). Additionally, HuR cleavage has been reported to function as a molecular switch that alters HuR affinity for target transcripts and promotes apoptosis progression (von Roretz et al. 2013a). In the present study, cleaved caspase-3 was not detected at any time point following APAP exposure (Supplementary Figure 1). Because hepatocyte death after APAP overdose is well established to occur predominantly through mitochondrial dysfunction and necrotic cell death, rather than caspase-3-dependent apoptosis (Jaeschke et al. 2018; Lambrecht et al. 2024), these findings indicate that APAP-induced liver injury is associated with HuR protein modification and cleavage through caspase-3 independent mechanisms. Accordingly, further studies are warranted to define the molecular pathways responsible for HuR cleavage and modification during APAP-induced hepatotoxicity.

### Heightened liver injury severity in *HuR*^Hep-/-^ mice at early time points following APAP overdose

We next assessed the progression of APAP-induced liver injury in WT and *HuR*^Hep-/-^ mice. Liver tissues and plasma were collected at 2, 6, and 24 hours after APAP overdose. Quantitative PCR confirmed the efficient *HuR* depletion in *HuR*^Hep-/-^ livers (Figure 2A). As expected, plasma ALT levels and hepatic necrotic area increased over time following APAP administration in both genotypes (Figure 2B-D). Compared with WT mice, *HuR*^Hep-/-^ mice exhibited significantly higher plasma ALT levels at both 2 and 6 hours post-overdose, along with more extensive liver necrosis at 6 hours. In contrast, the severity of liver injury was comparative between genotypes at 24 hours post overdose (Figure 2B-D). Overall, these data indicate that hepatocyte *HuR* deficiency is associated with increased early liver injury following APAP overdose (200 mg/kg), without a detectable difference in injury severity at later time points.

**Figure 2.**
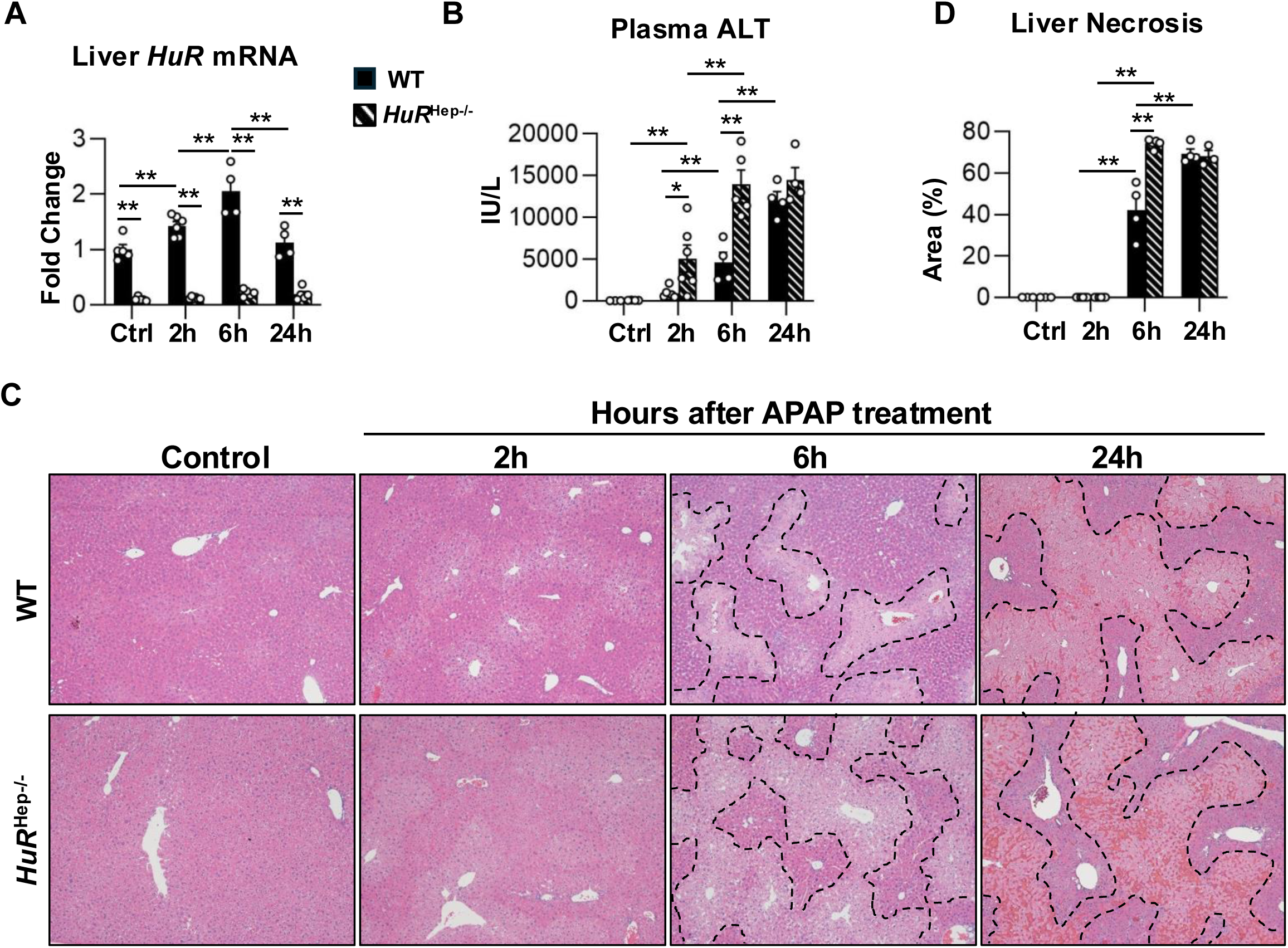
Liver injury is exacerbated in male *HuR*^Hep-/-^ mice after APAP overdose. HuR expression and liver injury were assessed at indicated time points following 200 mg/kg APAP. (A) qPCR analysis of liver HuR mRNA expression. (B) Plasma alanine aminotransferase (ALT) following APAP overdose. (C) Representative images of H&E-stained liver sections with dashed lines demarcating necrotic area. (D) Quantification of liver necrosis area outlined in C. Data are represented as mean ± SEM, n=3-6 mice per group. * p<0.05, ** p<0.01. Abbreviations: APAP, acetaminophen; H&E, hematoxylin and eosin; HuRHep-/-, hepatocyte-specific HuR knockout; WT, wild type.

### FTIR analysis reveals distinct biochemical signatures in WT and *HuR*^Hep-/-^ livers before and after 2-hour APAP exposure

FTIR spectroscopy was applied to liver sections from control and 2h APAP-treated WT and *HuR*^Hep-/-^ mice to characterize subtle biochemical differences associated with *HuR* deficiency and early APAP-induced injury. Representative FTIR spectra from three regions of interest (ROIs) located near central vein regions per sample and three samples per group (n=9 ROIs per group) were analyzed and compared (Supplementary Figure 2). Spectral changes were observed across three regions: (a) 1480 – 1420 cm⁻¹, corresponding primarily to lipids; (b) 1350 – 1200 cm⁻¹, associated with the amide III band of protein and nucleic acids; and (c) 1150 – 950 cm⁻¹, reflecting contributions from glycogen, nucleic acids, and phospholipid (Figure 3A). Changes were assessed based on peak height/intensity and peak shifts, which reflect quantity and structural alterations of these molecules.

**Figure 3.**
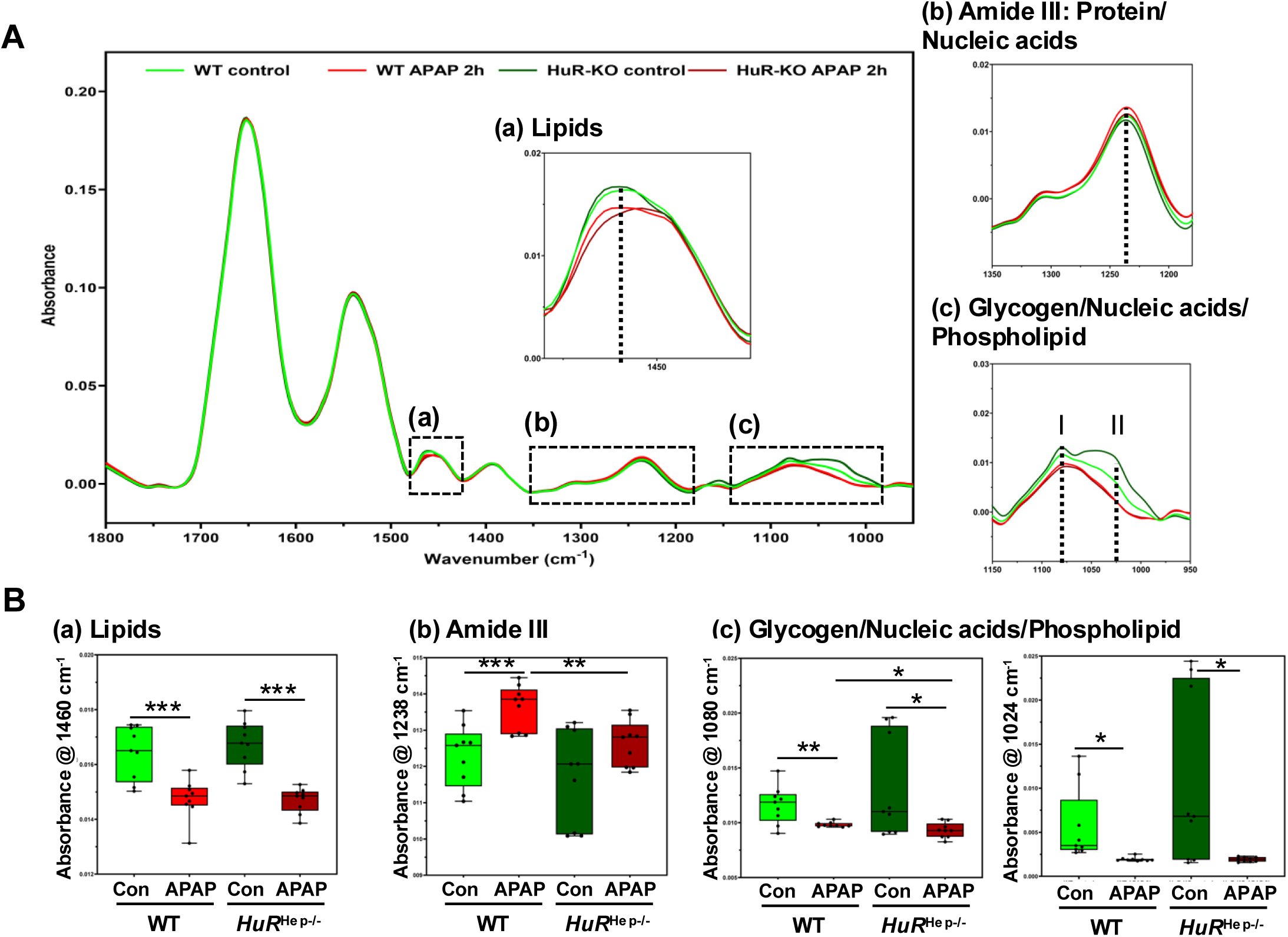
FTIR analysis reveals distinct biochemical signatures in WT and *HuR*^Hep-/-^ livers before and after 2-hour APAP exposure. Representative FTIR spectra were obtained from three regions of interest (ROIs) located near the central vein per sample, with three samples per group (n = 9 ROIs per group), and analyzed for comparison. (A) FTIR spectral plots highlighting characteristic wavenumber regions corresponding to (a) lipids, (b) amide III (proteins/nucleic acids), and (c) glycogen/nucleic acids/phospholipids. (B) Box plots quantifying the average maximum intensities at (a) 1460 cm⁻¹, (b) 1238 cm⁻¹, and (c) 1080 cm⁻¹ (left) and 1024 cm⁻¹ (right) across treatment groups. Data are represented as mean ± SEM, n=3 mice per group. * p<0.05, ** p<0.01, *** p<0.001. Abbreviations: FTIR, Fourier transform infrared; APAP, acetaminophen; HuRHep-/-, hepatocyte-specific HuR knockout; WT, wild type.

In the lipid-associated 1480 – 1420 cm⁻¹ region, both WT and *HuR*^Hep-/-^ mice exhibited significantly reduced peak intensity at 1460 cm⁻¹ following 2 hours of APAP treatment compared with their respective controls (p = 0.0007 for WT; p = 0.00001 for *HuR*^Hep-/-^) (Figure 3B, Left). The *HuR*^Hep-/-^ APAP 2h group also displayed a band shift toward lower wavenumbers compared to other groups, suggesting APAP-induced structural alterations in lipid molecules that are accentuated by *HuR* deficiency (Figure 3A). Within the 1350 – 1200 cm⁻¹ spectral region, the WT APAP 2h group exhibited a significant increase in peak intensity at 1238 cm⁻¹ compared with untreated WT controls (p = 0.001). In contrast, this APAP-induced increase was absent in *HuR*^Hep-/-^ livers, resulting in a significantly lower peak intensity in *HuR*^Hep-/-^ APAP 2h samples compared with WT APAP 2h (p = 0.003) (Figure 3B, Middle). In the 1150 – 950 cm⁻¹ spectral region, both APAP-treated WT and *HuR*^Hep-/-^ livers exhibited reduced peak intensities at 1080 cm^-1^ (p = 0.006 and p = 0.03, respectively) and 1024 cm^-1^ (p = 0.02 for both), suggesting reduced hepatic glycogen content after APAP treatment (Figure 3B, Right). Importantly, the 1080 cm^-1^ peak was further decreased in *HuR*^Hep-/-^ APAP 2h samples relative to WT APAP 2h (p = 0.05), suggesting that *HuR* loss modifies the hepatic metabolic response to APAP injury. Additionally, the *HuR*^Hep-/-^ control group exhibited large inter-sample variations in both the 1350 – 1200 cm⁻¹ and 1150 – 950 cm⁻¹ spectral regions (Figure 3B), indicating baseline metabolic heterogeneity in *HuR*-deficient livers. In contrast, no significant differences between WT and *HuR*^Hep-/-^ groups were detected in the amide I (1700 – 1600 cm⁻¹) and amide II (1570 – 1500 cm⁻¹) regions (Figure 3A). In summary, FTIR spectroscopy revealed significant, region-specific biochemical alterations in liver tissue as early as 2 hours after APAP exposure in both WT and *HuR*^Hep-/-^ mice, with *HuR* deficiency exacerbating or modifying lipid- and glycogen-associated spectral changes during early hepatotoxic injury.

### Altered APAP metabolite disposition in *HuR*^Hep-/-^ mice 2 hours after APAP overdose

Efficient early detoxification and elimination of reactive APAP metabolites are critical for limiting liver injury following APAP overdose (McGill and Jaeschke 2013). Bulk liver RNA-seq analysis of untreated WT and *HuR*^Hep-/-^ livers revealed dysregulation of multiple metabolic pathways in *HuR*^Hep-/-^ mice, including pathways associated with xenobiotic metabolism (Fig. 4A). More specifically, several significantly repressed DEGs identified in *HuR*^Hep-/-^ livers encoded glutathione S-transferases (*Gsts*), including *Gsta1*, *Gsta2*, *Gsta4*, *Gstm6*, and *Gstm7* (Figure 4B). These cytosolic, phase II detoxification enzymes catalyze the conjugation of glutathione (GSH) to electrophilic metabolites such as the reactive APAP metabolite NAPQI, as well as the glutathionylation of redox-sensitive proteins (Coles et al. 1988; Mazari et al. 2023). Although GSH conjugation predominates at therapeutic APAP doses, GST-mediated detoxification becomes increasingly important as hepatic GSH levels decline during overdose (Coles et al. 1988).

**Figure 4.**
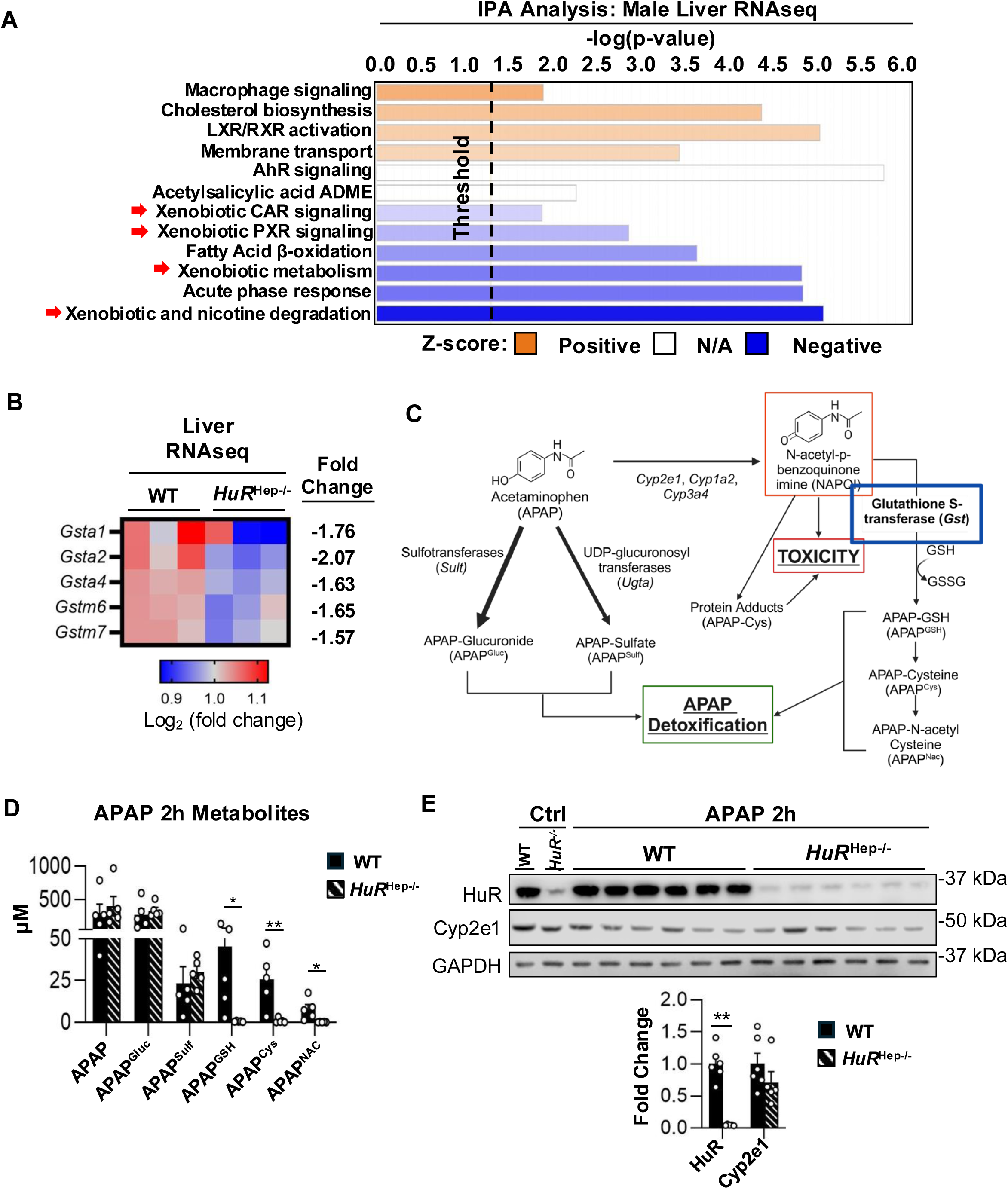
Dysregulation of APAP metabolism in *HuR*^Hep-/-^ mice 2 hours after APAP overdose. (A) IPA-based canonical pathway analysis of liver RNA-seq from untreated WT and *HuR*^Hep-/-^ male mouse livers; significantly dysregulated pathways had a -log(p-value) >1.3; Z >0 = pathway activation, Z<0 = pathway inhibition. (B) Heatmap of significantly repressed glutathione S-transferases (Gst) identified in the WT vs *HuR*^Hep-/-^ RNA-seq comparison. (C) Schematic of hepatic APAP metabolism. (D) LC-MS/MS analysis of APAP metabolites in plasma collected 2 hours following 200 mg/kg APAP treatment. (E) Western blot analysis of HuR and Cyp2e1 protein expression. Band intensity was quantified and normalized to GAPDH. Data are represented as mean ± SEM, n=3-6 mice per group. * p<0.05, ** p<0.01. Abbreviations: RNA-seq, RNA sequencing; IPA, Ingenuity Pathway Analysis; Cyp2e1, cytochrome P450 2e1; GAPDH, glyceraldehyde-3-phosphate dehydrogenase.

In both mice and humans, APAP undergoes extensive hepatic metabolism, and plasma levels of APAP and its downstream metabolites reflect hepatic metabolic capacity (Fischer et al. 1981; Prescott 1980) (Figure 4C). To evaluate APAP metabolism in *HuR*^Hep-/-^ mouse, APAP metabolites were quantified in plasma collected from WT and *HuR*^Hep-/-^ mice 2 hours after 200 mg/kg APAP overdose. Plasma levels of unconjugated APAP and the major phase II metabolites APAP-glucuronide (APAP^Gluc^) and APAP-sulfate (APAP^Sulf^) were comparable between WT and *HuR*^Hep-/-^ mice (Figure 4D). In contrast, consistent with reduced GSTs expression, plasma levels of downstream products of GSH conjugation to NAPQI, including APAP-GSH (APAP^GSH^), APAP-cysteine (APAP^Cys^), and APAP N-acetylcysteine (APAP^Nac^), were dramatically decreased in *HuR*^Hep-/-^ mice relative to WT controls. Specifically, at 2 hours after APAP overdose, plasma from *HuR*^Hep-/-^ mice contained 87-fold, 29-fold, and 78-fold lower levels of APAP^GSH^, APAP^Cys^, and APAP^Nac^, respectively (Figure 4D). Because GSH conjugation to NAPQI occurs only after phase I bioactivation of APAP (McGill and Jaeschke 2013), expression of the primary cytochrome P450 enzyme responsible for NAPQI formation, Cyp2e1, was also examined. However, Cyp2e1 protein levels did not differ between WT and *HuR*^Hep-/-^ mice (Figure 4E). Collectively, these findings indicate that altered APAP metabolite disposition in *HuR*-deficient mice is unlikely to result from differences in APAP bioactivation via Cyp2e1. Rather, these findings suggest impaired downstream detoxification, particularly reduced GSH conjugation, which may increase the availability of reactive NAPQI for covalent binding to hepatic proteins, a well-established contributor to APAP-induced liver injury.

### Aberrant mitochondrial structure and function in *HuR*^Hep-/-^ livers before and after APAP exposure

Mitochondrial dysfunction is central to the progression of APAP-induced hepatotoxicity (Ramachandran and Jaeschke 2019b). Mechanistically, APAP overdose impairs mitochondrial respiratory function in hepatocytes (Burcham and Harman 1991), while inherent defects in the electron transport chain (ETC) function predispose mitochondria to injury (Vercellino and Sazanov 2022). To assess mitochondrial respiration, we used an O2k fluorometer to measure respiration and H_2_O_2_ production of mitochondria isolated from untreated WT and *HuR*^Hep-/-^ male livers, as well as from WT and *HuR*^Hep-/-^ livers collected 2 hours after 200 mg/kg APAP treatment. Various electron transport chain substrates (pyruvate, ADP, glutamate, malate) and the uncoupler, FCCP, were added to mitochondria within the fluorometric chamber to assess mitochondrial respiratory capacity. Overall, mitochondrial respiration increased with the addition of each substrate and reached maximal levels following FCCP addition in mitochondria isolated from both WT and *HuR*^Hep-/-^ mice (Figure 5A). However, leak (basal) respiration and succinate-driven State 3 (State 3S) respiration were decreased significantly in mitochondria from untreated *HuR*^Hep-/-^ livers relative to WT controls (Figure 5A). In addition, mitochondrial redox status (H_2_O_2_^Basal^/O_2_^Basal^) was elevated in mitochondria isolated from untreated *HuR*^Hep-/-^ livers relative to WT, although OXPHOS capacity (Resp.^ADP^-Resp.^Basal^) did not differ between genotypes (Figure 5B). Following APAP treatment, mitochondrial respiration was impaired in both WT and *HuR*^Hep-/-^ mice, as indicated by markedly reduced OXPHOS capacity and increased redox potential in mitochondria isolated 2 hours after APAP overdose (Figure 5B). Because proper assembly and function of the electron transport chain (ETC) are essential for mitochondrial respiration and limiting oxidant stress (Vercellino and Sazanov 2022), we next assessed ETC protein expression. Western blot analysis revealed no differences in the expression of representative proteins from ETC complexes I-V between WT and *HuR*^Hep-/-^ livers under either control or APAP-treated conditions (Supplementary Figure 3A). Collectively, these findings indicate that impaired mitochondrial respiration in untreated *HuR*^Hep-/-^livers occurs independently of changes in ETC protein abundance and may predispose mitochondria to increased oxidant stress.

**Figure 5.**
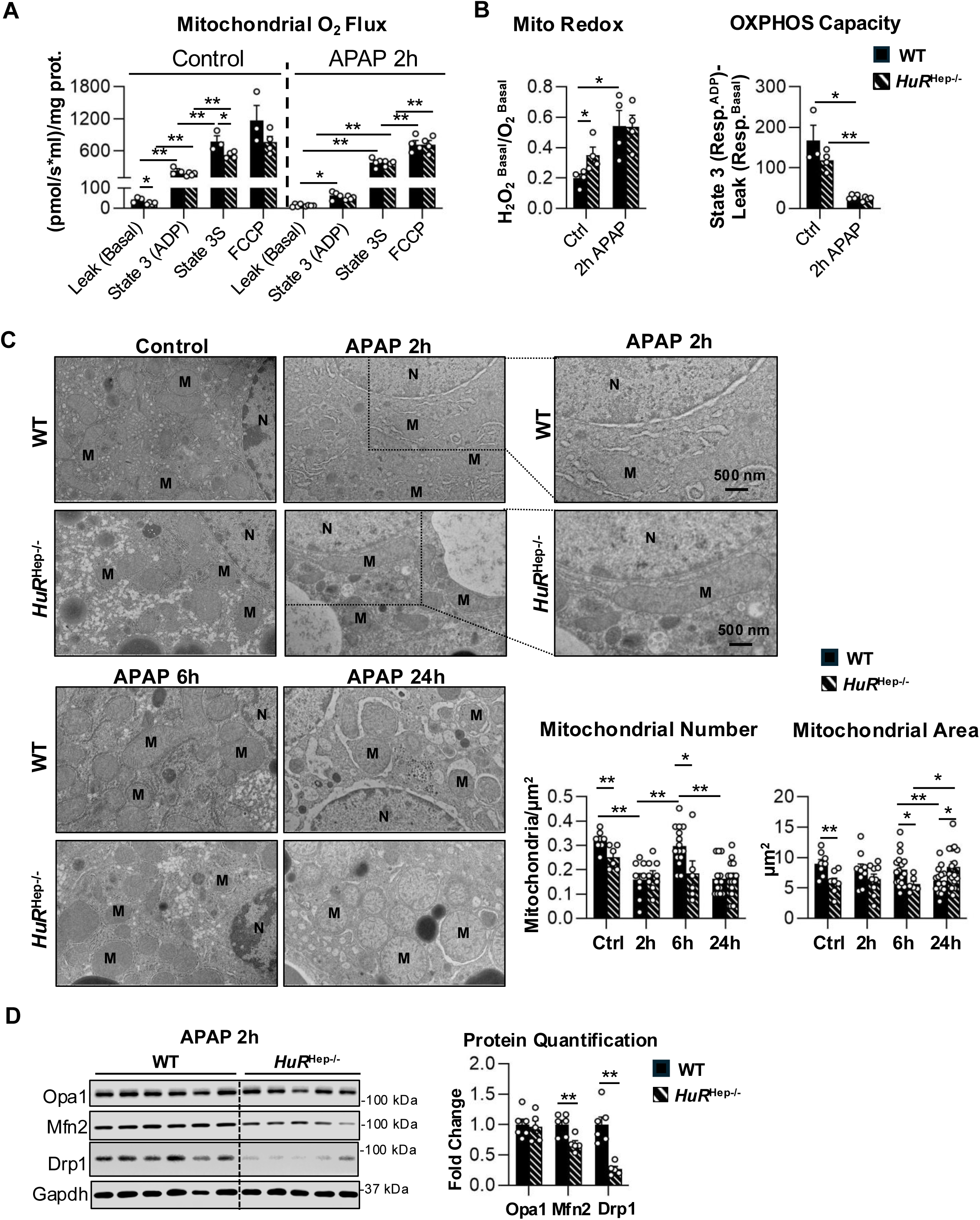
Mitochondrial respiration, morphology, and dynamics are disrupted in *HuR*^Hep-/-^livers before and after APAP exposure. (A) Mitochondrial respiration and H₂O₂ production were measured using an Oroboros O2k fluorometer in mitochondria isolated from WT and *HuR*^Hep-/-^livers under untreated (control) conditions and at 2 hours after APAP treatment (2 h APAP). Oxygen consumption was assessed sequentially following the addition of malate, ADP, succinate, and FCCP. (B) Mitochondrial redox status, expressed as basal H₂O₂ production normalized to basal O₂ flux, and OXPHOS capacity, calculated as State 3 respiration minus leak respiration. (C) Representative transmission electron micrographs (TEMs) of WT and *HuR*^Hep-/-^ mouse livers collected under control conditions and at indicated time points following APAP treatment. Mitochondrial number and area were quantified using ImageJ. (D) Western blot analysis of mitochondrial dynamics proteins Opa1, Mfn2, and Drp1 in livers collected 2 hours after APAP treatment. Protein levels were normalized to GAPDH. Data are represented as mean ± SEM, n=3-6 mice per group. * p<0.05, ** p<0.01. Abbreviations: ADP, adenine diphosphate; FCCP, carbonyl cyanide-p-(trifluoromethyoxy) phenylhydrazone; M, mitochondria; N, nucleus, Opa1, optic atrophy 1; Mfn2, mitofusion 2; Drp1, dynamin-related protein 1.

Defects in mitochondrial structure are closely linked to mitochondrial dysfunction under conditions of cellular stress (Eisner et al. 2018; Picard et al. 2013). To assess mitochondrial morphology, transmission electron microscopy (TEM) was used to examine hepatocyte mitochondria from untreated livers and from livers collected 2, 6, and 24 hours following APAP overdose. Overall, mitochondrial number and morphology in hepatocytes changed dynamically in response to APAP exposure and hepatocyte-specific *HuR* deficiency (Figure 5C). In untreated conditions, hepatocytes from *HuR*^Hep-/-^ livers contained fewer mitochondria and reduced total mitochondrial area compared with WT controls (Figure 5C). Despite this reduction, mitochondria in *HuR*^Hep-/-^ hepatocytes displayed a more circular morphology, as quantified by the circularity index (4π × area/perimeter²; ImageJ) (Figure 5C and Supplementary Figure 3B). At 2 hours following APAP overdose, mitochondrial number was decreased in both WT and *HuR*^Hep-/-^hepatocytes relative to untreated controls; however, mitochondria at this time point exhibited a more elongated morphology (Figure 5C). Importantly, extremely large and elongated mitochondria were observed specifically in *HuR*^Hep-/-^ livers at 2 hours post-APAP exposure (Figure 5C, insert). By 6 hours post-overdose, mitochondrial number increased in WT hepatocytes relative to the 2-hour time point, whereas this increase was not observed in *HuR*^Hep-/-^ livers (Figure 5C). As a result, *HuR*^Hep-/-^ hepatocytes exhibited reduced mitochondrial area compared with WT at this time point (Figure 5C). At 24 hours, mitochondrial circularity and surface area in WT hepatocytes returned to levels comparable to untreated controls. In contrast, *HuR*^Hep-/-^ hepatocytes maintained significantly higher mitochondrial circularity and surface area relative to WT (Figure 5C and Supplementary Figure 3B). Taken together, these data demonstrate that both APAP exposure and hepatocyte-specific *HuR* deficiency disrupt mitochondrial morphology. In particular, the dynamic changes in mitochondrial number observed in WT livers, characterized by an early decrease at 2 hours, followed by partial recovery at 6 hours and a subsequent decline at 24 hours, were blunted in *HuR*^Hep-/-^ livers, suggesting impaired mitochondrial remodeling in response to APAP-induced stress.

Canonical mitochondrial fusion and fission proteins play essential roles in large-scale remodeling of mitochondrial structure and function (Picard et al. 2013). Accordingly, we next assessed the expression of key mitochondrial fusion and fission proteins 2 hours after APAP overdose, a time point associated with pronounced morphological alterations. At this time point, *HuR*^Hep-/-^ livers exhibited a 1.5-fold reduction in the outer mitochondrial membrane fusion protein mitofusin 2 (Mfn2) and a 2.4-fold reduction in the cytoplasmic mitochondrial fission protein dynamin-related protein 1 (Drp1) compared with WT livers (Figure 5D). Collectively, the coordinated reduction in Mfn2 and Drp1 expression is consistent with the aberrant mitochondrial morphology observed in *HuR*^Hep-/-^ hepatocytes and provides a mechanistic link between disrupted mitochondrial dynamics and the impaired respiratory capacity observed in *HuR*-deficient livers. Together, these findings indicate that hepatocyte *HuR* deficiency compromises mitochondrial dynamics, thereby exacerbating both structural remodeling and functional mitochondrial deficits under basal conditions and following APAP-induced liver injury.

### Increased mitochondrial protein release, oxidant stress, and cell death in *HuR*^Hep-/-^ livers 2 hours after APAP overdose

Defects in mitochondrial respiration and morphology contribute to APAP-induced hepatotoxicity by promoting mitochondrial oxidant stress and release of mitochondrial proteins into the cytosol (Picard et al. 2013). Accordingly, markers of mitochondrial oxidant stress and mitochondrial dysfunction were assessed in WT and *HuR*^Hep-/-^ mouse livers at 2 hours following APAP overdose (Figure 6). Phosphorylation and mitochondrial translocation of c-JUN N-terminal kinase (pJNK) are established indicators of JNK activation and increased mitochondrial ROS production (Ramachandran and Jaeschke 2019a). At 2 hours post-overdose, the pJNK/JNK ratio was increased 1.6-fold in the cytoplasmic fraction of *HuR*^Hep-/-^ male mouse livers relative to WT (Figure 6A). In contrast, mitochondrial translocation of pJNK and total JNK did not differ between WT and *HuR*^Hep-/-^ mouse livers at this time point (Figure 6B), suggesting that JNK activation was unlikely to be a major contributor to the heightened liver injury observed in *HuR*^Hep-/-^ mice at this time point.

**Figure 6.**
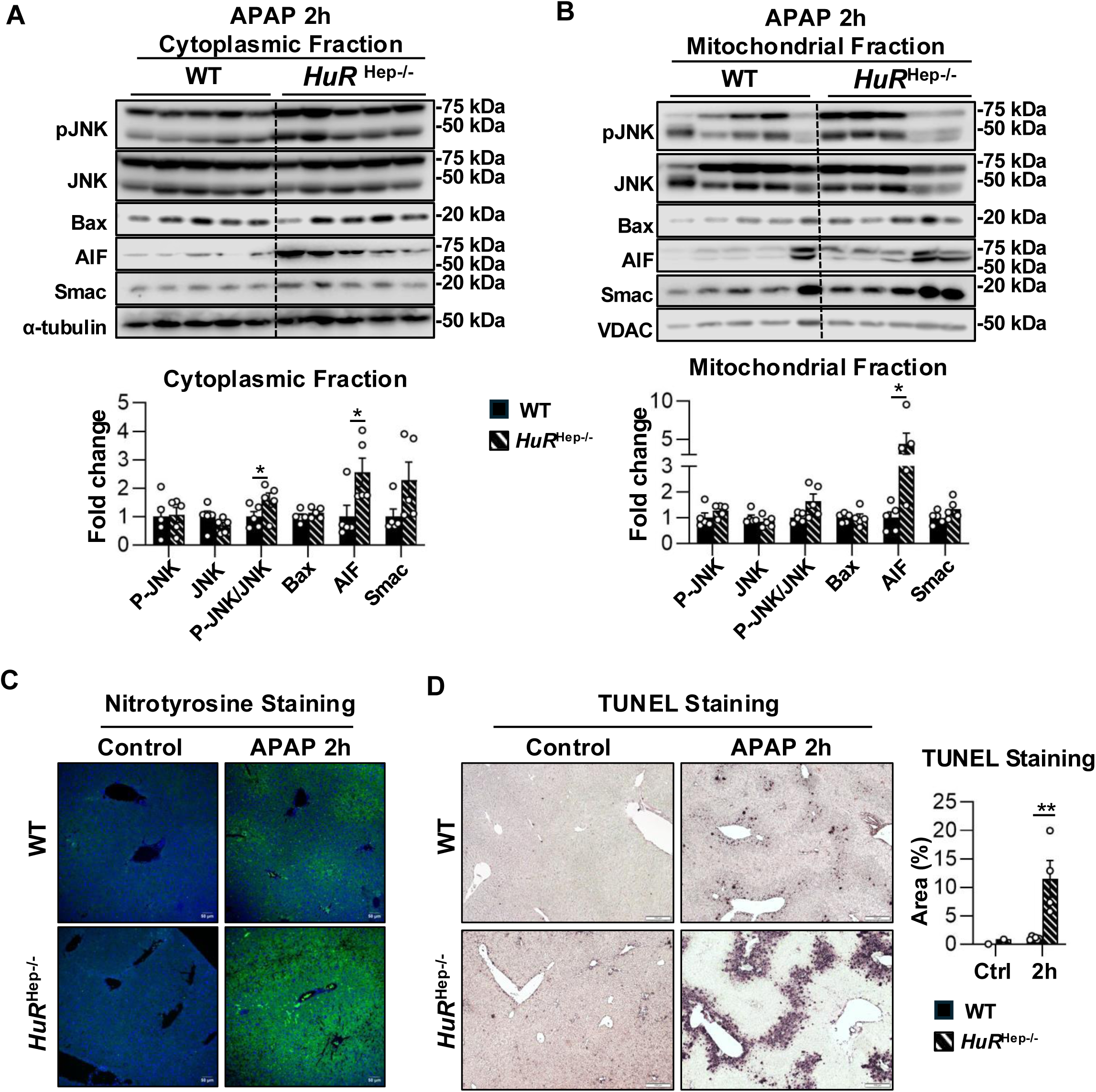
Increased mitochondrial oxidant stress and mitochondrial protein release in APAP-treated *HuR*^Hep-/-^ male mice 2 hours after APAP overdose. (A-B) Western blot analysis of proteins isolated from cytoplasmic (A) and mitochondrial (B) liver fractions collected 2 hours after APAP overdose. Protein levels were quantified densitometrically and normalized to VDAC (mitochondrial) or α-tubulin (cytosolic). (C) Representative immunofluorescence images of nitrotyrosine staining in liver sections collected 2 hours after APAP overdose. (D) Representative images of TUNEL staining in liver sections collected 2 hours after APAP overdose, positive staining area quantified using Fiji (image J). Data are represented as mean ± SEM, n=3-5 mice per group. * p<0.05, ** p<0.01. Abbreviations: pJNK, phosphorylated c-Jun N terminal kinase; AIF, apoptosis-inducing factor; VDAC, voltage-dependent anion-selective channel 1; TUNEL, Terminal deoxynucleotidyl transferase dUTP nick end labeling.

To more directly assess mitochondrial oxidant stress, immunofluorescence staining for nitrotyrosine protein adducts was performed. Following APAP overdose, nitrotyrosine adduct formation is largely restricted to mitochondria damaged by ROS, making nitrotyrosine adducts a relatively specific marker of mitochondrial oxidant stress (Cover et al. 2005). Consistent with enhanced mitochondrial oxidative stress, immunofluorescence staining revealed visibly higher levels of nitrotyrosine adducts in *HuR*^Hep-/-^ male mouse livers compared with WT at 2 hours post-overdose (Figure 6C). In parallel, levels of the mitochondrial intermembrane space protein apoptosis-inducing factor (AIF), a marker of severe mitochondrial dysfunction (Bajt et al. 2006; Masubuchi et al. 2005), were markedly increased in both mitochondrial and cytoplasmic fractions of *HuR*^Hep-/-^ male mouse livers relative to WT (Figure 6A and B). Specifically, AIF levels were increased 2.6-fold in mitochondrial fractions and 4.4-fold in cytoplasmic fractions of *HuR*^Hep-/-^ livers at 2 hours following APAP overdose. Following release into the cytoplasm, AIF translocates to the nucleus, where it functions as an endonuclease that cleaves DNA and promotes hepatocyte necrosis (Bajt et al. 2006). Consistent with increased mitochondrial AIF release and indicative of DNA damage, terminal deoxynucleotidyl transferase dUTP nick-end labeling (TUNEL) staining was increased approximately 13-fold in *HuR*^Hep-/-^ mouse livers compared with WT at 2 hours post-overdose (Figure 6D). Taken together, these findings suggest that enhanced mitochondrial dysfunction and oxidative stress are key drivers of early hepatocyte necrosis in *HuR*^Hep-/-^ male mouse livers following APAP overdose.

### Persistent mitochondrial dysfunction, enhanced APAP-protein adduct accumulation, and impaired glutathione recovery exacerbate hepatotoxicity in *HuR*-deficient livers at 6 hours after APAP overdose

More severe liver injury persisted in *HuR*^Hep-/-^ male mouse livers at 6 hours following APAP overdose (Figure 2). Accordingly, canonical markers of mitochondrial dysfunction and APAP-induced hepatotoxicity were assessed in WT and *HuR*^Hep-/-^ livers at this later time point. As observed at 2 hours, cytoplasmic JNK activation was modestly increased in *HuR*^Hep-/-^ male mouse livers but did not reach statistical significance, and overall JNK activation and mitochondrial translocation were comparable between WT and *HuR*^Hep-/-^ livers at 6 hours (Figure 7A and B). In contrast, cytoplasmic levels of the mitochondrial intermembrane proteins AIF and Smac were robustly increased in *HuR*^Hep-/-^ male mouse livers compared to WT (Figure 7A). Specifically, AIF and Smac levels were increased 7.1-fold and 3.6-fold, respectively, in cytoplasmic fractions of *HuR*^Hep-/-^ livers at 6 hours post-overdose (Figure 7A).

**Figure 7.**
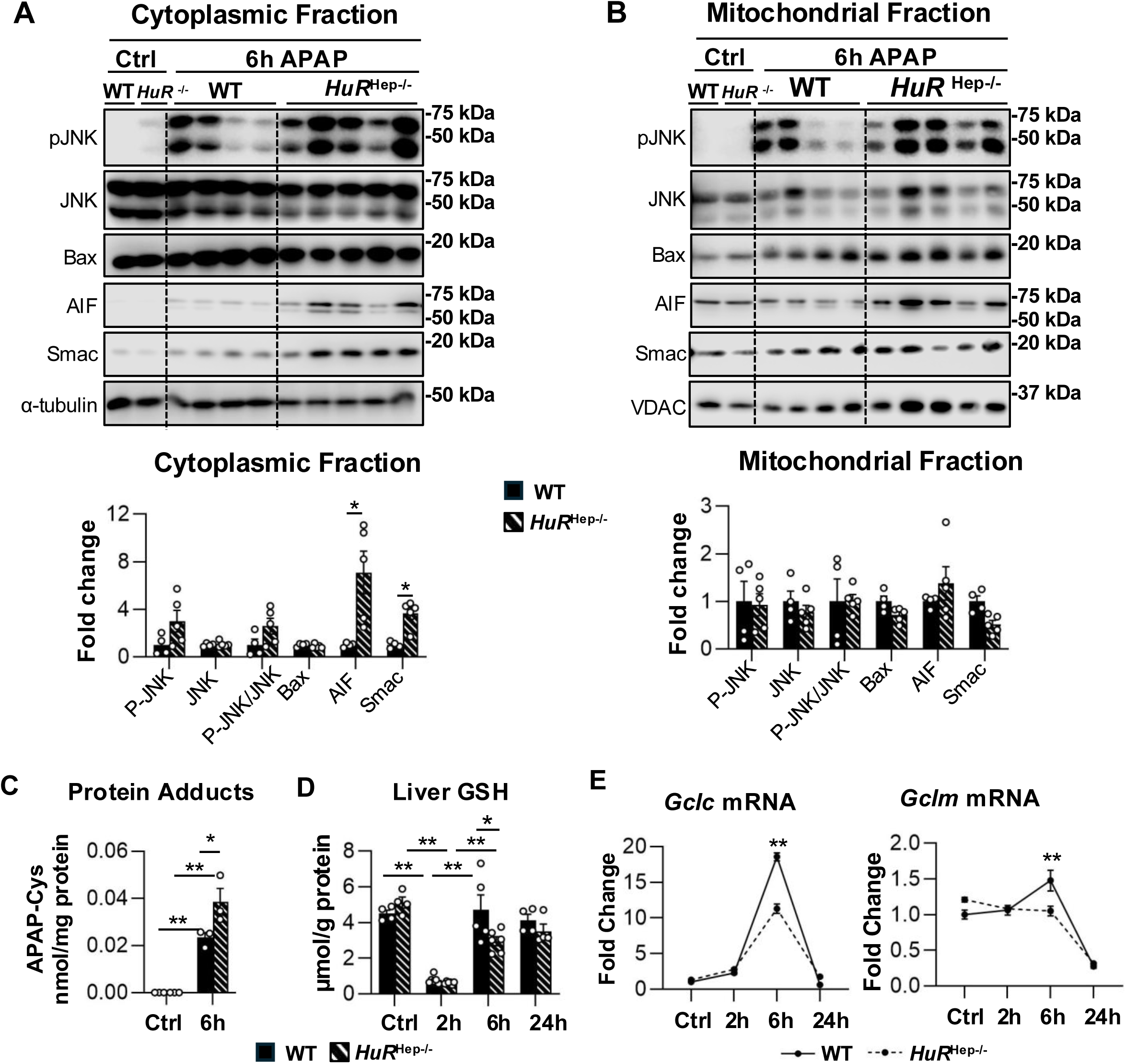
Persistent mitochondrial dysfunction, enhanced APAP-protein adduct accumulation, and impaired glutathione recovery in *HuR*^Hep-/-^ livers 6 hours after APAP overdose. (A-B) Western blot analysis of proteins isolated from cytoplasmic (A) and mitochondrial (B) liver fractions collected 6 hours after APAP overdose. Protein levels were quantified densitometrically and normalized to α-tubulin (cytosolic) or VDAC (mitochondrial). (C) High performance liquid chromatography (HPLC) quantification of APAP-protein adducts (APAP-Cys) in WT and *HuR*^Hep-/-^ livers at 6 hours after APAP overdose. (D) Total hepatic glutathione (GSH) levels measured using a modified Tietze method at indicated time points following APAP overdose. Total GSH includes both reduced GSH and oxidized glutathione (GSSG). (E) qPCR analysis of liver *Gclc* and *Gclm* mRNA expression over time following APAP overdose. Data are represented as mean ± SEM, n=3-5 mice per group. * p<0.05, ** p<0.01. Abbreviations: pJNK, phosphorylated c-Jun N-terminal kinase; AIF, apoptosis-inducing factor; VDAC, voltage-dependent anion-selective channel 1; GSH; glutathione; Gclc, glutamate cysteine ligase catalytic subunit; Gclm, glutamate cysteine ligase modifier subunit.

Altered APAP metabolite disposition in *HuR*-deficient mice (Figure 4D) may impair downstream detoxification, thereby increasing the availability of reactive NAPQI for covalent binding to hepatic proteins and formation of APAP-Cys protein adducts, a well-established contributor to APAP-induced liver injury (Jaeschke and Ramachandran 2024). Consistent with this, APAP-Cys adduct levels were increased 1.6-fold in *HuR*^Hep-/-^ male mouse livers compared with WT at 6 hours post-overdose (Figure 7C). In parallel, total hepatic GSH levels were reduced 1.6-fold in *HuR*^Hep-/-^ male mouse livers relative to WT at the 6-hour time point (Figure 7D). Glutamate cysteine ligase (Gcl), the rate-limiting enzyme in GSH synthesis, consists of a catalytic (Gclc) and a modifier (Gclm) subunits (Lu 2013). Both *Gclc* and *Gclm* mRNA levels were robustly induced in WT livers 6 hours after APAP overdose (Figure 7E). In contrast, induction of *Gclc* and *Gclm* was blunted in *HuR*^Hep-/-^ livers by approximately 1.6-fold and 1.4-fold, respectively (Figure 7E), which explains the delayed recovery of hepatic GSH levels at 6 hours (Figure 7D). Taken together, these findings suggest that increased accumulation of APAP-Cys protein adducts, accompanied by impaired GSH recovery, potentially due to reduced induction of Gclc and Gclm, contributes to the persistent liver injury observed in *HuR*^Hep-/-^ male mice at 6 hours following APAP overdose.

### Loss of HuR enhances pro-inflammatory gene expression and impairs Cyclin D1 induction following APAP overdose

Sterile inflammation − mainly driven by the activity of macrophages and neutrophils − plays a crucial role in resolving liver injury following APAP overdose (Jaeschke and Ramachandran 2020; Rodrigues et al. 2023). Immunohistochemistry (IHC) staining for the macrophage marker, F4/80, and the neutrophil marker, lymphocyte antigen 6 family member G (Ly6g), revealed similar immune cell abundance in WT and *HuR*^Hep-/-^ livers following APAP overdose (Figure 8A). However, qPCR analysis showed increased expression of *Ly6g* and pro-inflammatory cytokines, including C-C motif chemokine ligand 2 (*Ccl2*), tumor necrosis factor α (*Tnfa*), and interleukin 6 (*IL6*), in *HuR*^Hep-/-^ male mouse livers compared to WT, especially at the 24-hour timepoint following APAP overdose (Figure 8B). These findings suggest that *HuR* loss promotes a more pro-inflammatory environment compared to WT during APAP overdose-induced hepatotoxicity.

**Figure 8.**
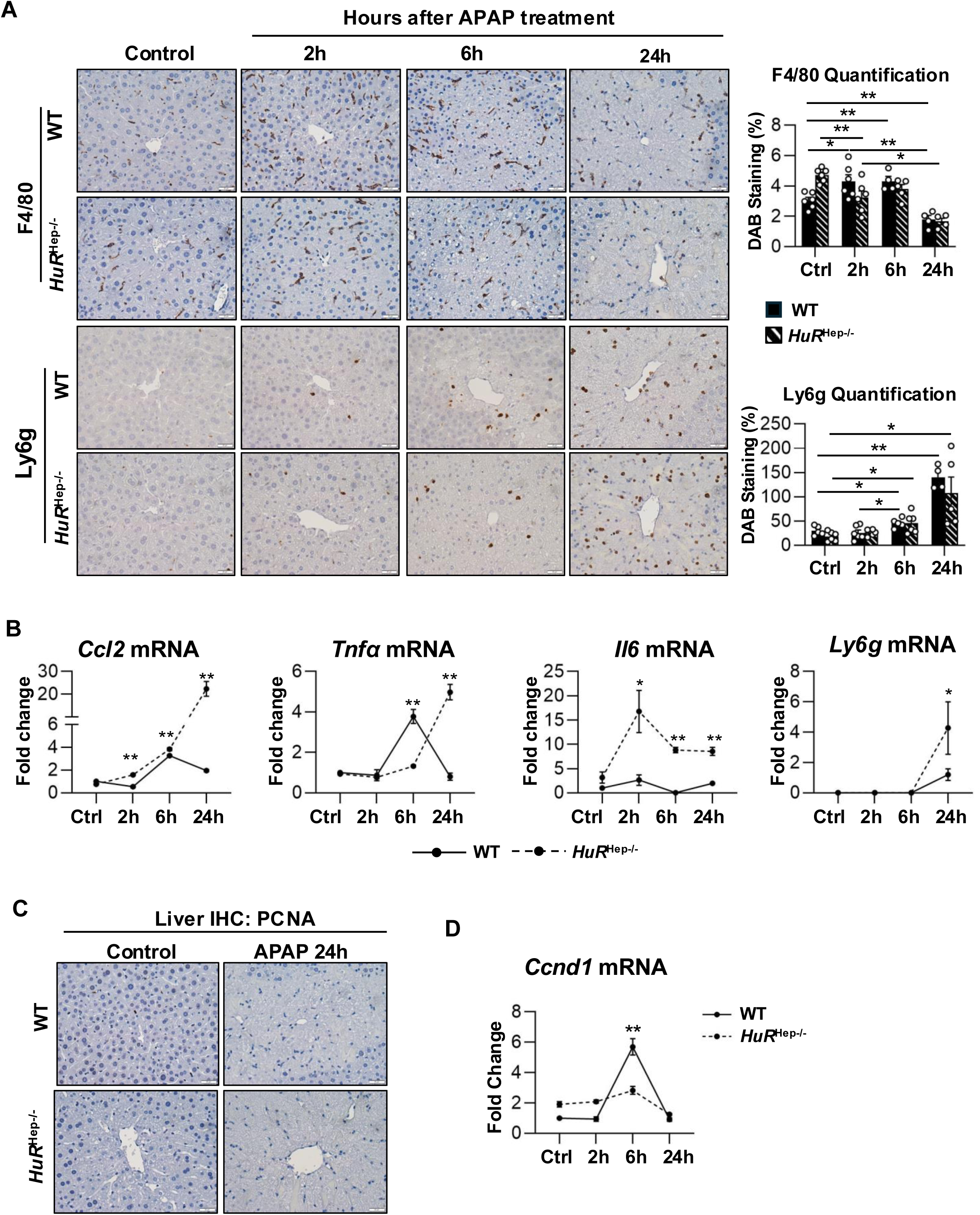
Loss of HuR enhances pro-inflammatory gene expression and impairs *Cyclin d1* induction following APAP-overdose. (A) Representative IHC images of liver sections stained for F4/80 (top) and Ly6g (bottom). Percent positive staining area and neutrophil number were quantified using Image J. (B) qPCR analysis of hepatic *Ccl2*, *Tnfa*, *Il6*, and *Ly6g* mRNA expression. (C) Representative IHC images of liver sections stained for PCNA. (D) qPCR analysis of hepatic *Ccnd1* mRNA expression at indicated time points following APAP overdose. The mRNA levels were normalized to *Hprt1*. Data are represented as mean ± SEM, n=3-5 mice per group. * p<0.05, ** p<0.01. Abbreviations: IHC, Immunohistochemistry; HPF, high power field; Ly6g, lymphocyte antigen 6; DAB, 3,3’-diaminobenzidine; Ccl2, C-C motif chemokine ligand 2; Tnfa, tumor necrosis factor alpha; Il6, interleukin 6; PCNA, proliferating cell nuclear antigen; Ccnd1, cyclin d1.

Hepatocytes proliferation is critical for liver regeneration after APAP overdose (Bhushan and Apte 2019). To assess hepatocyte proliferation, IHC staining for proliferating cell nuclear antigen (PCNA) was performed on liver tissues from WT and *HuR*^Hep-/-^ male mice on a C57BL/6N background. Although liver regeneration is less well characterized in the C57BL/6N substrain, studies in C57BL/6J mice have reported the increase of PCNA-positive hepatocytes at 24 hours following a moderate APAP dose (300 mg/kg), whereas a severe dose (600 mg/kg) delays PCNA induction in male livers (Bhushan et al. 2014). In the present model, PCNA-positive hepatocytes were not detected in either WT or *HuR*^Hep-/-^ male mouse livers at 24 hours following a 200 mg/kg APAP overdose compared with untreated controls (Figure 8C). Given that C57BL/6N mice are significantly more susceptible to APAP-induced hepatotoxicity than C57BL/6J mice (Duan et al. 2016), these data suggest that 200 mg/kg APAP induces a severe injury in C57BL/6N male mice, sufficient to delay the onset of liver regeneration. Although PCNA-positive cells were not detected at this dose, Cyclin D1, a key regulator of hepatocyte cell cycle entry (Albrecht and Hansen 1999), was significantly induced at the mRNA level in WT male livers 6 hours after APAP overdose; in contrast, this induction was absent in *HuR*^Hep-/-^ livers (Figure 8D). Taken together, these findings indicate that liver regeneration was delayed in both WT and *HuR*^Hep-/-^ male mice at 24 hours following a 200 mg/kg APAP overdose, and that Cyclin D1 expression and thus early cell cycle entry may be further impaired in *HuR*^Hep-/-^ livers relative to WT.

## DISCUSSION

HuR is a ubiquitously expressed RNA-binding protein that regulates cellular survival and homeostasis by modulating the stability and processing of numerous target transcripts (Eppler et al. 2025; Srikantan and Gorospe 2012). The present study comprehensively examined the role of hepatocyte HuR in APAP-induced hepatotoxicity across the metabolism/early injury (2 hours), injury (6 hours), and early recovery (24 hours) phases following APAP overdose. After APAP exposure, increased HuR mRNA expression, HuR protein cleavage, and the appearance of higher-molecular weight HuR species closely tracked with the progression of liver injury. During the metabolism and injury phases, hepatocyte-specific *HuR* deficiency exacerbated liver injury. Specifically, enhanced liver injury during the early metabolism phase was associated with altered oxidative APAP metabolite disposition, increased nitrosative stress, and mitochondrial dysfunction. At 6 hours post-overdose, more severe liver injury in *HuR*^Hep-/-^ mice was accompanied by increased APAP-protein adduct formation, delayed GSH recovery, and elevated mitochondrial protein release. Finally, although impaired hepatocyte proliferation at 24 hours in both WT and *HuR*^Hep-/-^ mice reflected the overall severity of injury at this dose, *HuR*^Hep-/-^ livers exhibited distinct increases in pro-inflammatory cytokine expression and impaired induction of Cyclin D1. Collectively, these findings identify HuR as a critical regulator of mitochondrial integrity and liver homeostasis during APAP-induced hepatotoxicity (Figure 9).

**Figure 9.**
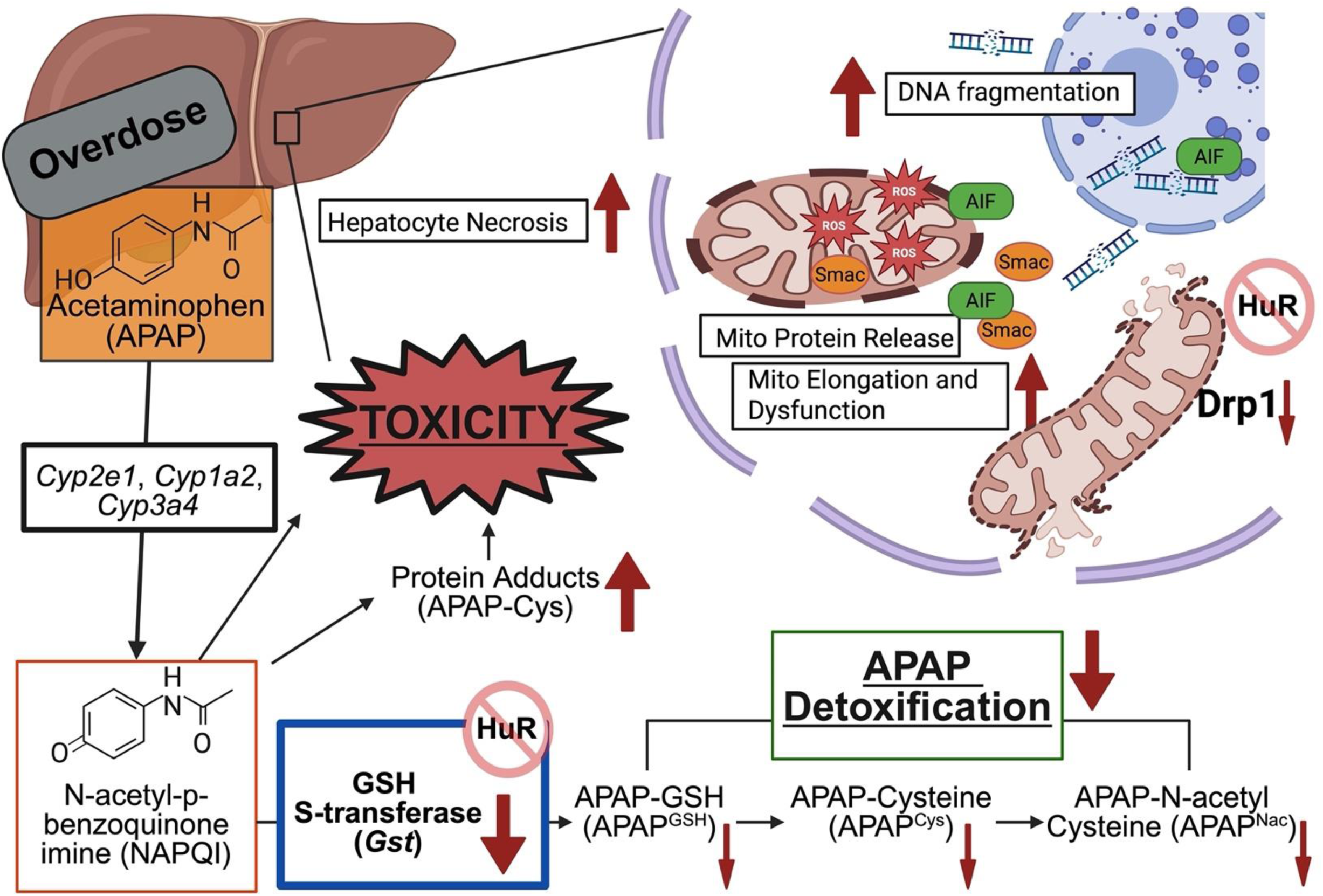
Hepatocyte-specific *HuR* deficiency exacerbates APAP-induced hepatotoxicity in male mice. Schematic depicting how loss of hepatocyte HuR amplifies APAP-induced liver injury through coordinated defects in xenobiotic metabolism and mitochondrial homeostasis. HuR deficiency reduces glutathione S-transferase (Gst) expression and disrupts APAP detoxification, leading to increased NAPQI burden and enhanced APAP-protein adduct formation. Concomitantly, impaired induction of glutathione synthesis delays GSH recovery, further limiting detoxification capacity. HuR loss also dysregulates mitochondrial dynamics by reducing Drp1 and Mfn2 expression, resulting in abnormal mitochondrial morphology, compromised respiratory function, and increased mitochondrial membrane permeabilization. These defects promote mitochondrial protein release and nuclear DNA fragmentation, thereby intensifying hepatocyte necrosis and overall liver injury in *HuR*^Hep-/-^ male mice. Abbreviations: NAPQI, N-acetyl-p-benzoquinone imine; Gst, glutathione S-transferase; GSH, glutathione; Mfn2, mitofusion 2; Drp1, dynamin-related protein 1.

An important finding of this study is that induction of hepatic HuR expression, together with its cleavage and post-translational modification, closely parallels the time course of liver injury following APAP overdose. Cellular stress, including oxidative stress, is known to induce both HuR expression and HuR cleavage (Ashour et al. 2024; Banos-Jaime et al. 2024; Mazroui et al. 2008; Talwar et al. 2011; von Roretz and Gallouzi 2010; von Roretz et al. 2013b). In addition, HuR dimerization at the RNA recognition motif 1 (RRM1) interface can be mediated by redox-sensitive disulfide bond formation (Benoit et al. 2010), suggesting that oxidative conditions such as those induced by APAP overdose may favor HuR multimerization. In the present study, both HuR cleavage products and higher-molecular weight HuR species were most prominent at 24 hours, coinciding with peak liver injury. Although the in vivo functions of these HuR modifications remain incompletely understood, HuR cleavage has been shown to promote apoptotic cell death in HeLa cells under lethal stress (Mazroui et al. 2008), whereas HuR also functions as a pro-survival factor in many cancers (Wu and Xu 2022). This apparent functional plasticity is thought to be regulated, at least in part, by post-translational modifications such as protein cleavage and protein-protein interactions (Grammatikakis et al. 2017). For example, HuR dimerization and higher-order complex formation enhance its affinity for target transcripts (Pabis et al. 2019), while HuR cleavage is required for muscle cell differentiation through stabilization of differentiation-related mRNAs (Beauchamp et al. 2010). Despite evidence that HuR cleavage alters target specificity, the mechanisms and functional consequences of HuR modification in APAP-injured livers remain unclear. The strong correlation between HuR modifications and serum ALT levels supports a close association between HuR modification and hepatocellular injury severity. Moreover, the appearance of both HuR cleavage products and higher-molecular weight species in the absence of caspase-3 activation suggests that HuR undergoes caspase-independent modification during APAP-induced hepatotoxicity, consistent with the predominance of necrotic cell death in this model. Further studies are therefore warranted to define the molecular pathways responsible for HuR cleavage and modification during APAP-induced hepatotoxicity.

Another import finding of the present study is the identification of a role for HuR in regulating APAP metabolism. Previous studies have identified a metabolism-associated function for HuR in maintaining liver homeostasis (Eppler et al. 2025). For example, liver ribonucleoprotein immunoprecipitation (RNP-IP) combined with bulk RNA sequencing revealed dysregulation of multiple metabolic pathways in *HuR*-deficient mouse livers and direct HuR binding to transcripts encoding key enzymes involved in endobiotic and xenobiotic metabolism (Subramanian et al. 2022). Consistent with these findings, our study revealed marked dysregulation of GSH-dependent APAP metabolism in *HuR*^Hep-/-^ mouse livers. Specifically, plasma levels of GSH-conjugated APAP metabolite levels, including APAP^GSH^, APAP^Cys^, and APAP^Nac^, were markedly reduced in *HuR*^Hep-/-^ mice 2 hours after APAP overdose, whereas levels of the major non-GSH-dependent metabolites, APAP^Gluc^ and APAP^Sulf^, were comparable between genotypes. In parallel, RNA-seq analysis revealed reduced mRNA expression of multiple Gsts, including *Gsta1*, *Gsta2*, *Gsta4*, *Gstm6*, and *Gstm7*, in *HuR*^Hep-/-^ mouse livers. Although evidence supporting a dominant role for Gsts in APAP metabolism remains limited, the contribution of specific Gst isoforms to APAP-induced hepatotoxicity has been partially explored. For instance, *Gstp1/2*- and *Gstm*-deficient mice are resistant to APAP-induced hepatotoxicity despite exhibiting largely intact APAP metabolism (Arakawa et al. 2012; Henderson et al. 2000). In human cohorts, loss-of-function variants in *GSTT1* have been associated with improved outcomes following APAP overdose (Buchard et al. 2012). While these observations suggest that inhibition of certain Gst isoforms may be protective in specific contexts, substantial variability exists in Gst substrate specificity (Eaton and Bammler 1999). Moreover, although members of the Gstα family can catalyze GSH conjugation to NAPQI *in vitro* (Coles et al. 1988), their in vivo role in APAP-induced hepatotoxicity remains poorly defined. Future studies will therefore focus on characterizing Gst isoform expression and enzymatic activity following APAP overdose and on quantifying APAP metabolites in the liver and bile of *HuR*^Hep-/-^ mice to more precisely define the contribution of HuR-dependent metabolic regulation to APAP detoxification.

In the present study, hepatocyte HuR emerged as a dynamic and multifaceted regulator of mitochondrial biology. Enhanced liver injury in *HuR*-deficient livers following APAP overdose was associated with pronounced alterations in mitochondrial morphology, dysregulated mitochondrial dynamics, impaired respiratory function, and increased mitochondrial permeabilization. Previous studies have highlighted diverse roles for HuR in regulating mitochondrial biology. For example, liver-specific *HuR* deficiency increases susceptibility to diet-induced MASLD by reducing expression of electron transport chain components, including cytochrome c (*Cycs*), NADH:ubiquinone oxidoreductase subunit B6 (*Ndufb6*), and ubiquinol–cytochrome c reductase binding protein (*Uqcrb*) (Zhang et al. 2020). In HEK293 cells, HuR maintains mitochondrial DNA integrity by facilitating cytoplasmic translocation of the long noncoding RNA, RNA processing endoribonuclease (*RMRP*) (Cascajo et al. 2016), while in differentiating B cells, mitochondrial energy production depends on HuR-mediated pre-mRNA processing of dihydrolipoamide S-succinyltransferase (*Dlst*) (Diaz-Munoz et al. 2015). HuR has also been shown to protect against mitochondrial oxidant stress through regulation of stress-responsive genes such as heme oxygenase 1 (Dery et al. 2020). Consistent with these reports, *HuR*-deficient livers in the current study exhibited reduced expression of both the mitochondrial fission protein Drp1 and the fusion protein Mfn2 following APAP overdose, accompanied by the appearance of large, elongated mitochondria at the 2-hour time point. These findings are supported by prior in vitro work demonstrating that HuR binds the 3′ untranslated region of *Drp1* mRNA and that HuR loss promotes mitochondrial elongation (Bae et al. 2019). Although whether HuR regulates Mfn2 directly or indirectly remains to be determined, *HuR*-deficient hepatocytes from both untreated and APAP-exposed livers contained mitochondria with a more circular morphology. Increased mitochondrial circularity is commonly associated with loss of mitochondrial membrane potential and mitochondrial dysfunction following APAP exposure (Umbaugh et al. 2021). Reflecting cumulative defects in mitochondrial structure and function, release of mitochondrial intermembrane space proteins was markedly increased in *HuR*^Hep-/-^ livers at both 2 and 6 hours after APAP overdose. Because release of mitochondrial endonucleases is a key driver of nuclear DNA fragmentation and hepatocyte necrosis (Bajt et al. 2006), these findings implicate mitochondrial dysfunction as a central contributor to the heightened liver injury observed in *HuR*^Hep-/-^ mice. In addition, HuR has been reported to regulate mitophagy through upregulation of PARKIN and BNIP3L (Yu et al. 2021), and inhibition of autophagy has been observed in male *HuR*-deficient livers following APAP overdose (Lu et al. 2024). Given that autophagic clearance of damaged mitochondria is a critical protective mechanism in APAP-induced liver injury (Chao et al. 2018; Ni et al. 2012), and that large, swollen mitochondria accumulated in *HuR*^Hep-/-^hepatocytes at 24 hours post-overdose, future studies should determine whether HuR directly regulates mitophagy during APAP-induced hepatotoxicity.

Compensatory liver regeneration, characterized by peri-necrotic hepatocyte proliferation, is essential for recovery following APAP overdose (Bhushan and Apte 2019). In an incremental dose model using C57BL/6J male mice, a moderate APAP dose (300 mg/kg) induces robust liver regeneration at 24 hours, whereas a higher dose (600 mg/kg) results in severe liver injury accompanied by delayed regeneration, cell cycle arrest, and reduced Cyclin D1 expression (Bhushan et al. 2014). In the present study, neither WT nor *HuR*^Hep-/-^ male mouse livers exhibited PCNA-positive staining at 24 hours following treatment with 200 mg/kg APAP. Given the extensive hepatic necrosis and markedly elevated serum ALT levels observed in both genotypes at this time point, it is likely that 200 mg/kg APAP represents a non-regenerative dose in C57BL/6N male mice. Although the use of a non-regenerative APAP dose limited direct assessment of liver regeneration, our findings nonetheless identify multiple mechanisms, including dysregulated APAP metabolism and mitochondrial dysfunction, through which hepatocyte-specific *HuR* deficiency exacerbates APAP-induced hepatotoxicity in male mice.

## FUNDING

This work was supported by the National Institutes of Health (P30 GM118247, R01DK119131) and the University of Kansas Medical Center (Lied grant, Bridging Award).

## DECLARATION OF CONFLICTING INTERESTS

The authors declared no potential conflicts of interest regarding the research, authorship, and/or publication of this article.

---

**Supplementary table S1:**
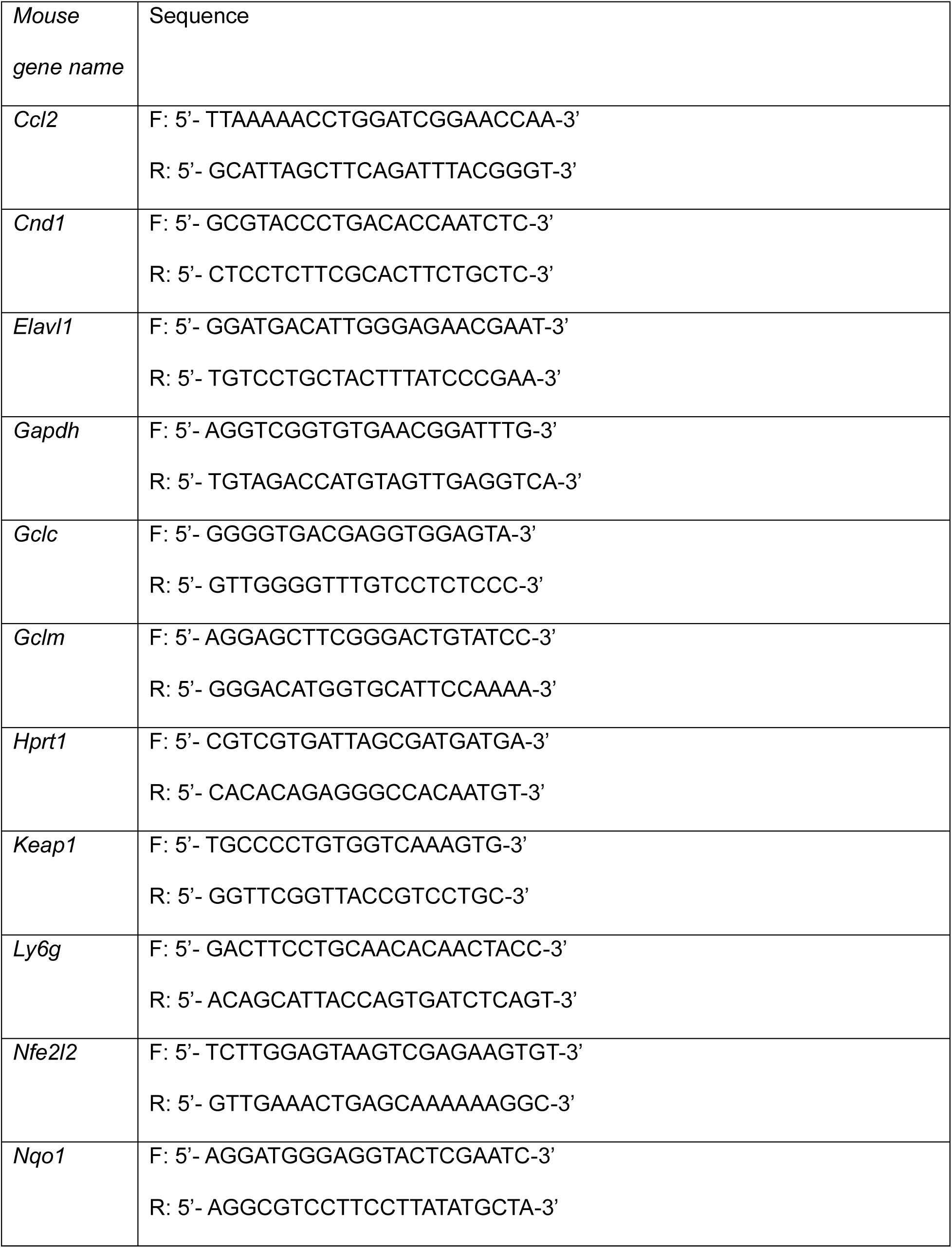
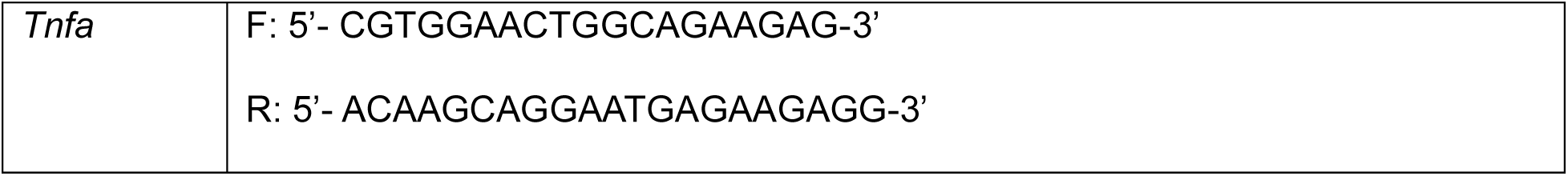

**Supplementary Figure 1.**
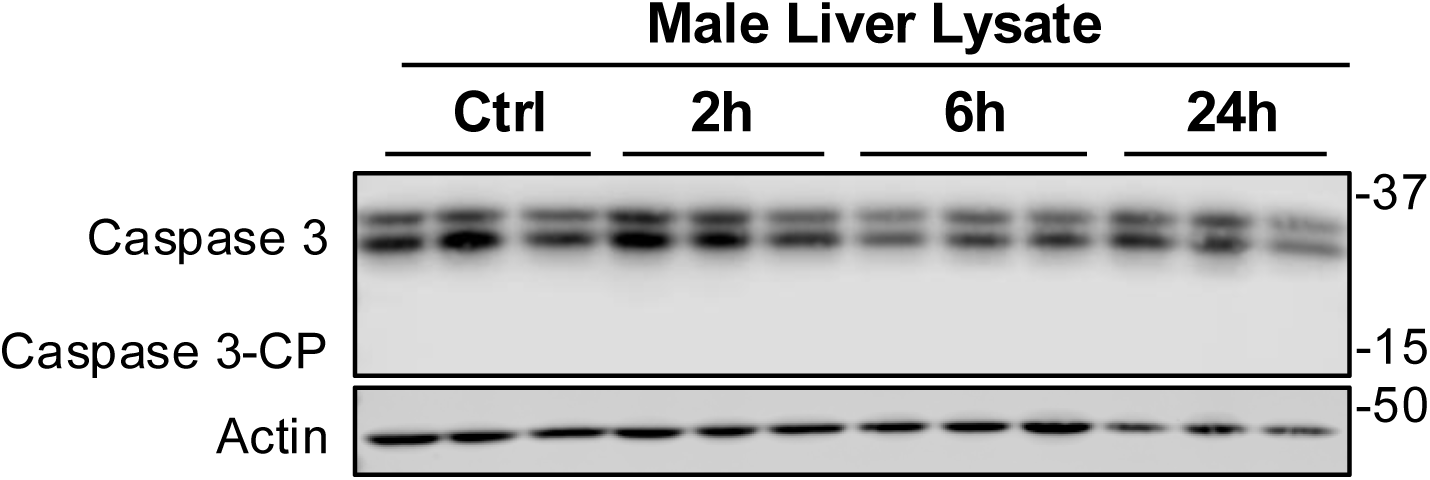
Caspase 3 cleavage was not detected in mouse livers following APAP-overdose. Proteins isolated from male livers collected at indicated time points following 200 mg/kg APAP were analyzed by Western blot. Abbreviations: Caspase 3-CP, Caspase3 cleavage product.

**Supplementary Figure 2.**
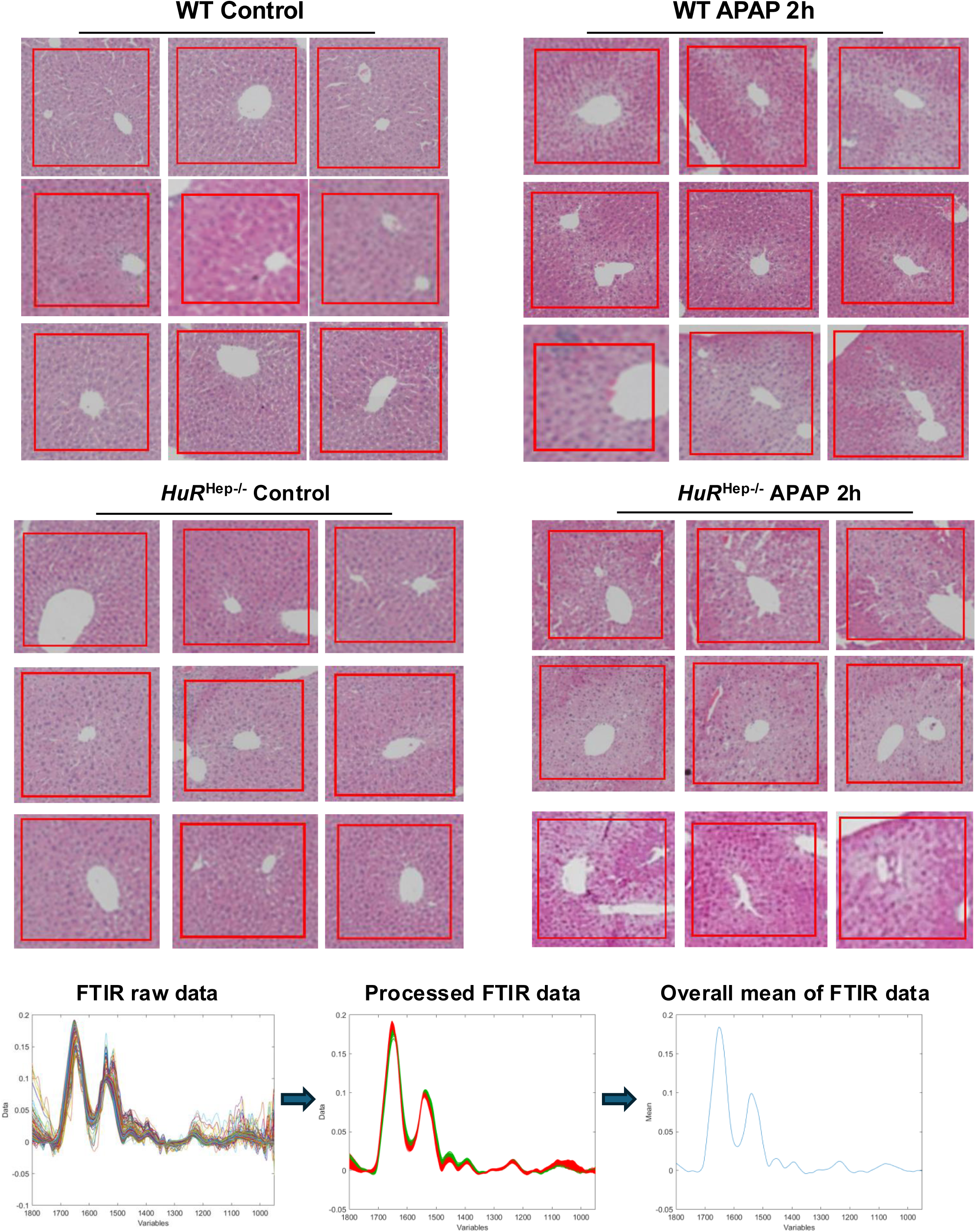
H&E-stained images of selected regions of interest (ROIs) used for FTIR analysis. Three ROIs located near the central vein were selected per sample, with three samples per group (n = 9 ROIs per group) for spectral evaluation.

**Supplementary Figure 3.**
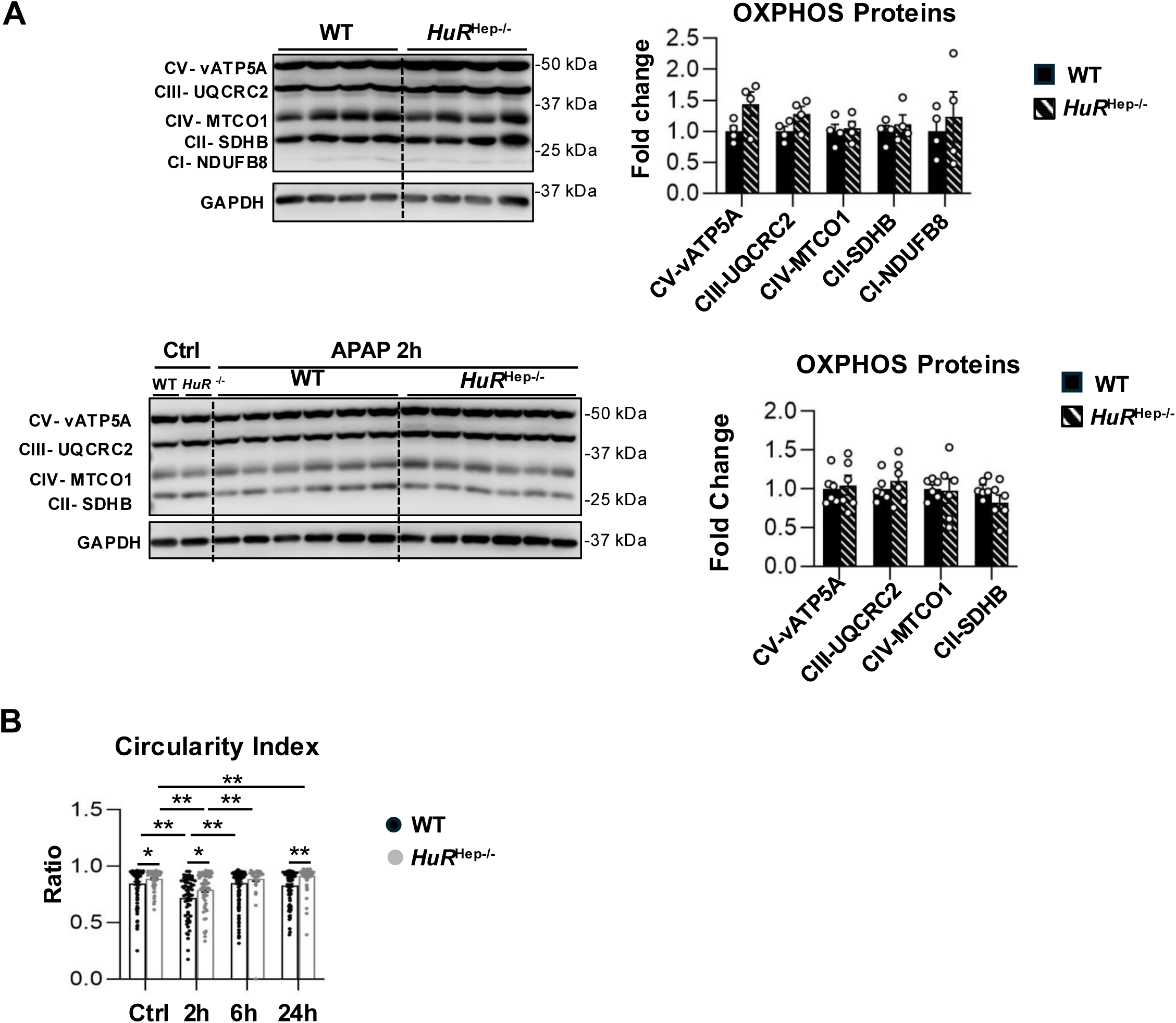
Hepatocyte-specific *HuR* knockout does not alter electron transport chain (ETC) complex I-V protein expression but affects mitochondrial morphology and size. (A) Western blot analysis of mitochondrial OXPHOS complex proteins isolated from livers of WT and *HuR*^Hep-/-^. Protein levels were normalized to GAPDH. Data are presented as mean ± SEM (n = 5 mice per group). *p < 0.05, **p < 0.01. (B) Quantification of mitochondrial circularity in livers from WT and *HuR*^Hep-/-^ under untreated conditions and at 2, 6, and 24 hours following 200 mg/kg APAP administration. Data are presented as mean ± SEM; each symbol represents an individual mitochondrion (n = 44–178). *p < 0.05, **p < 0.01.

## Notes

### Competing Interest Statement

The authors have declared no competing interest.

